# piRNA-like small RNAs target transposable elements in a Clade IV parasitic nematode

**DOI:** 10.1101/2022.02.21.481297

**Authors:** Mona Suleiman, Asuka Kohnosu, Ben Murcott, Mehmet Dayi, Rebecca Pawluk, Akemi Yoshida, Mark Viney, Taisei Kikuchi, Vicky L. Hunt

**Affiliations:** Department of Biology and Biochemistry, University of Bath, Bath, BA2 7AY UK; Parasitology, Department of Infectious Dieses, Faculty of Medicine, University of Miyazaki, Miyazaki, 889-1692 Japan; Laboratory of Genomics, Frontier Science Research Center, University of Miyazaki, Miyazaki, 889-1692 Japan; Department of Evolution, Ecology and Behaviour, University of Liverpool, Liverpool, L69 7ZB, UK

**Keywords:** PIWI pathway, piRNA, *Strongyloides*, small-interfering RNA, microRNA, small RNA, helminth, nematode, parasite, transposable elements

## Abstract

The small RNA (sRNA) pathways identified in the model organism *Caenorhabditis elegans* are not widely conserved across nematodes. For example, the PIWI pathway and PIWI-interacting RNAs (piRNAs) are involved in regulating and silencing transposable elements (TE) in most animals but have been lost in nematodes outside of the *Caenorhabditis elegans* group (Clade V), and little is known about how nematodes regulate TEs in the absence of the PIWI pathway. Here, we investigated the role of sRNAs in the Clade IV parasitic nematode *Strongyloides ratti* by comparing two genetically identical adult stages (the parasitic female and free-living female). We identified putative small-interfering RNAs, microRNAs and tRNA-derived sRNA fragments that are differentially expressed between the two adult stages. Two classes of sRNAs were predicted to regulate TE activity including (i) a parasite-associated class of 21-22 nt long sRNAs with a 5’ uracil (21-22Us) and monophosphate modification, and (ii) 27 nt long sRNAs with a 5’ guanine/adenine (27GAs) and polyphosphate modification. The 21-22Us show striking resemblance to the 21U PIWI-interacting RNAs found in *C. elegans*, including an AT rich upstream sequence, overlapping loci and physical clustering in the genome.

## Background

Small RNAs (sRNA) are short non-coding RNAs important for the regulation of gene expression via post-transcriptional gene silencing. They regulate the expression of at least 30% of genes in humans and are associated with chromatin structure, mRNA translation and the regulation of transposable element (TE) activity ^1,2,3^. Three main sRNA classes have been described in eukaryotes; microRNAs (miRNAs), small-interfering RNAs (siRNAs), and piwi-interacting RNA (piRNAs), classified based on their biogenesis, function and interaction with specific Argonaute proteins ^4^. The majority of sRNA research has been carried out in model organisms including *Drosophila melanogaster*, *Mus musculus* and *Caenorhabditis elegans*. *C. elegans* belongs to the Clade V nematodes and possess all three classes of sRNA like other model organisms. However, recent studies have shown that sRNA pathways are highly diverged in nematodes and *C. elegans* does not closely represent the sRNAs used by more distantly related nematodes, including parasitic species ^5^. For example, the PIWI pathway involved in the production of piRNAs is important in regulating TE activity and has been well characterised in *C. elegans* but has been lost in nematodes outside of the Clade V, including *Strongyloides* spp. ^675^. It is still not clear how nematodes outside of Clade V compensate for the loss of piRNAs to regulate TE activity.

piRNAs were first identified in *D*. *melanogaster* as PIWI-clade Argonaute interacting sRNAs and have subsequently been discovered in most other animals, including *M. musculus,* humans and *C. elegans* where they regulate TE activity, particularly in the germline ^8,9^. TEs are mobile DNA sequences that move around the genome from one location to another, inserting randomly and causing mutations ^8^. They play important roles in the evolution of eukaryotic organisms but can have detrimental effects to the genome and require tight regulation ^10,11^. Interestingly, while the role of piRNAs is widely conserved across eukaryotes including their role in fertility and protecting the germline from TEs ^12,13,14^, the PIWI pathway and piRNA biogenesis and mechanisms of action has diverged between organisms ^2^. For example, in *D*. *melanogaster* and *M*. *musculus*, piRNAs are often involved in the ping-pong cycle, are 24 - 30 nucleotides (nt) long with a bias for a 5’ uracil (5’ U) and they silence transposons through perfect antisense complementarity to their target sequences ^15,16^. This is not the case in *C. elegans* where piRNAs are 21nt long and although have a bias for a 5’ U, they don’t require perfect complementarity to their target sequence and are not involved in the ping-pong cycle. Instead, piRNAs in *C. elegans* initiate the synthesis of 22 long siRNAs with a 5’ guanine (22G) through RNA-dependent RNA polymerases (RdRPs) that silences complementary target sequences ^5,13,14^. The absence of piRNAs in nematodes outside of Clade V leads us to consider if an alternative sRNA pathway can compensate for the loss of piRNAs in nematodes outside of Clade V and regulate TE activity. It is therefore essential that we study nematodes outside of the *C. elegans* Clade including both parasitic and free-living nematodes, to better understand the diversity of sRNAs involved in regulating TEs and the role of sRNAs in parasitism.

Although both miRNAs and siRNAs pathways are found in all nematodes studied to date, their roles in gene regulation and mechanisms of silencing are still not fully understood. In contrast to piRNAs, miRNA sequences and the miRNA pathway show greater conservation among animals. miRNAs are a class of sRNA of 20 – 23 nt in length important for the regulation of protein-coding genes with diverse functions including the differentiation of larval stages and adult development ^17,18^. Complementarity of the miRNA and its target mRNA occurs through the seed sequence found in nucleotides 2 – 8 ^19,20^. More than 250 miRNAs have been identified in *C. elegans* where each of them can target more than one mRNA and each mRNA can be targeted by more than miRNA, increasing the complexity of gene regulation. The siRNAs, in comparison, are approximately 21 – 27 nt in length, and are important in chromatin regulation, transcriptional regulation, RNA degradation and protein modification ^21^. Classes of siRNAs can usually be classified by features such as their sequence length, 5’ starting nucleotide and 5′ end modifications ^7^, and the specific Argonaute protein they are loaded onto in a siRNA pathway ^4^. In *C. elegans*, processing of siRNAs by the enzyme Dicer creates primary siRNA with a 5’ monophosphate (5’pN). The primary siRNAs interact with specific Argonaute proteins, depending on their pathway, and create a complex with RdRPs, that uses the target transcript as a template for synthesis of the secondary siRNA, which are not Dicer-processed and typically have a 5’ triphosphate modification ^22^. In contrast to miRNAs that can target many mRNA through their small seed sequence, siRNAs are perfectly antisense to their specific target sites ^23^.

Here, we have investigated the role of sRNAs in the endogenous regulation of genes and TEs in the nematode *Strongyloides ratti,* a well-established laboratory model of nematode parasitism ^24^. *Strongyloides* species are gastrointestinal parasites which infect an estimated 600 million people globally causing chronic morbidity and, more rarely, fatal disseminated strongyloidiasis ^25^. They also infect animals causing substantial economic loss in livestock practices ^26^. The life cycle of *S. ratti* includes genetically identical free-living (FLF) and parasitic (PF) adult female stages. Direct comparison between these two adult life cycle stages can uncover genetic features associated with parasitism including differences in sRNA and TE activity. The *S. ratti* genome has been sequenced and assembled into a highly contiguous reference genome (two autosomes in single scaffolds and the X-chromosome in ten main scaffolds) ^27^ which enables an accurate genetic analysis of sRNAs and their targets. We have sequenced sRNAs that are expressed in PF and FLF *S. ratti*. The sRNAs were classified into classes or subsets of classes of sRNAs that were differentially expressed between PF and FLF life cycle stages. We identify two classes of sRNAs that are predicted to target TEs, that were differentially expressed between life cycle stages. The sRNAs associated with the parasitic stage shared multiple features in common with piRNAs including similar length (21-22 nt), a 5’ uracil, 5’ monophosphate modification, overlapping loci and an upstream AU-rich sequence; representing the first set of piRNA-like sRNAs outside of Clade V nematodes. We also identified miRNA families which are upregulated in the parasitic stage and tRNA fragments upregulated specifically in the free-living stage.

## Results

### *S. ratti* parasitic and free-living adult females express similar proportions of miRNAs and other sRNAs

sRNA expression in genetically identical PF and FLF *S. ratti* was investigated using two library types; (i) enriched for sRNAs with a 5’ monophosphate modification (5’pN enriched library), or (ii) RppH-treated to increase the cloning efficiency of 5’ polyphosphorylated and 5’ capped sRNAs (5’ modification-independent library). Reads were classified as either miRNAs, or as sRNAs derived from tRNAs (tRFs), rRNA (rsRNA), or as putative siRNAs originating from either protein-coding genes (including CDS and intronic regions), intergenic region or TEs. The most abundantly expressed class of sRNAs identified in the 5’pN enriched library was miRNAs with lengths of 21 - 23 nt, which made up 17.4% and 11.0% of total PF and FLF reads, respectively. The sRNAs originating from intergenic regions ranging in length between 21 – 24 nt were the second most highly expressed class of sRNA in both the PF and FLF (3.88% and 1.75% of all reads, respectively) (**Figure 1a & b, Supplementary Data 1**). Interestingly, 21-22 nt sRNAs originating from CDS and TEs were expressed at higher levels in the PF than the FLF (1.15% and 0.2% of 21-22nt reads in PF and FLF, respectively). In contrast, tRFs were more highly expressed in the FLF (0.7% and 1.6% in PF and FLF, respectively) (**Figure 1b, Supplementary Data 1**). Overall, there were more unique tRFs expressed in the FLF *cf.* PF (1036 and 185 sequences, respectively). tRFs were significantly upregulated (EdgeR, FDR < 0.01) in the FLF compared with the PF 5’pN-enriched libraries, and primarily originated from the central region of the mature tRNA sequences (**Supplementary Fig. 1**). The sRNA sequences expressed in the 5’pN-enriched library predominantly started with a 5’ uracil, consistent with the most common 5’ starting base for miRNAs (**Figure 1a & b)**.

**Figure 1:**
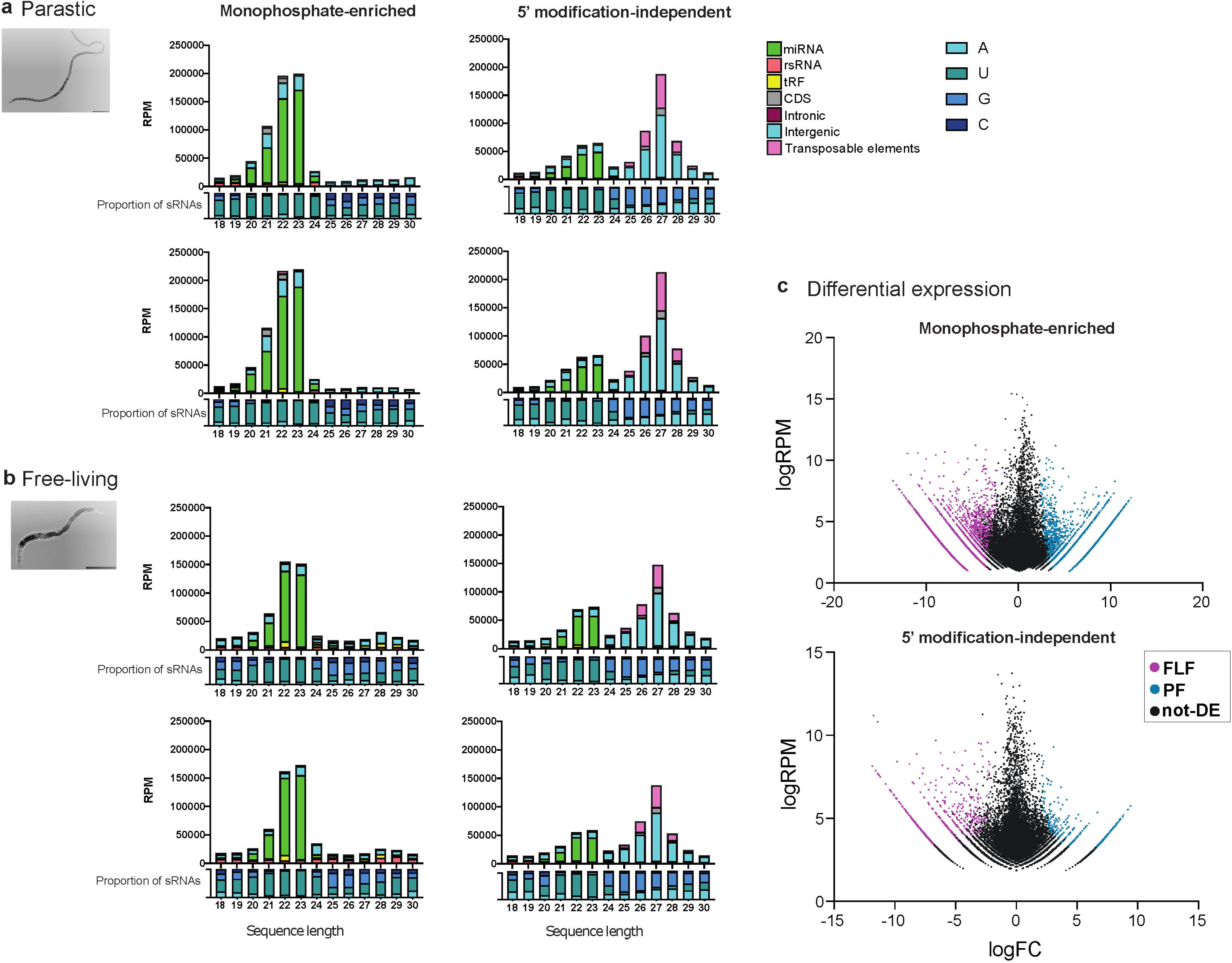
sRNA classification and differential expression. Classification of sRNAs expressed by parasitic female (PF) and free-living female (FLF) stages of *S. ratti* including sRNAs (a) enriched for a 5’ monophosphate modification, or (b) 5’ modification-independent sequences which includes sRNAs with a 5’ monophosphate or polyphosphate modifications. Graphs on the top show the classification of sRNAs as either miRNAs, rRNA-derived sRNAS (rsRNA), tRNA-derived sRNAs (tRFs) or as putative siRNAs originating from either protein-coding genes (CDS or intronic regions), intergenic regions or transposable elements (TE). RPM = reads per million. Graphs below the x-axis show the proportion of the first 5’ nucleotide for each length of sRNA. (c) Differential expression of sRNAs with a 5’ monophosphate modification (top), and 5’ modification-independent library (bottom). Significantly upregulated sequences are highlighted in pink (FLF) and blue (PF) (FDR of < 0.01, fold change of > 2) and sequences that are not differentially expressed are shown in black (logRPM = log reads per million, logFC = log fold change).

RppH treatment removes 5’ modifications in sRNA including 5’triphosphate and other 5’ modifications, thus sRNAs that are observed only in RppH-treated libraries are likely to have 5’ modifications. As expected, we observed similar peaks of 5’pN sRNAs including the miRNAs across both libraries (**Figure 1a & b)**. In addition, sRNAs between 24-30 nt in length were enriched in the 5’ modification-independent libraries indicating that *S. ratti* PF and FLF also express sRNAs with a 5’ modification. The 26-28 nt sRNAs originating from intergenic spaces and TE sequences were the most highly expressed class of sRNA in the 5’ modifications-independent libraries for both the PF and FLF. Together, intergenic- and TE-derived sRNAs comprised 65.9% and 41.2% of all sRNAs sequences in the 5’ modification-independent libraries for PF and FLF, respectively (**Figure 1a & b**). Overall, the sRNA expression profiles for FLF and PF in the 5’ modification-independent library were similar *i.e.* 27 nt sRNAs with a 5’ modification were most highly expressed, followed by 22-23 nt miRNAs with a 5’ monophosphate (**Supplementary Data 2**). In the 5’ modification-independent library, sRNA sequences between 18-23 in length, predominantly started with a 5’ uracil and were classified as miRNAs, and also identified in the 5’pN-enriched library.

sRNA sequences between 24-30 nt in length predominantly started with either a guanine or adenine at the 5’ end (**Figure 1a & b)**. To identify if specific sRNAs were upregulated in the PF *cf.* the FLF, a differential expression analysis was carried out for both libraries using edgeR ^28^. We found that 22.3% (n = 5584) and 21.9% (n = 5478) of 5’pN sRNA sequences (n = 25,047) were significantly upregulated in the PF and FLF life cycle stages, respectively. In the 5’ modification-independent library, 1.6% (n= 730) and 2.8% (n= 1278) of all sRNA sequences (n= 43,473) were significantly upregulated in the PF and FLF, respectively (**Figure 1c**) (FDR< 0.01, **Supplementary Data 3**). Together, these results indicate that distinct sets of sRNAs are upregulated in the PF and the FLF, suggesting they have specific roles in each life cycle stage. Interestingly, some sRNA sequences such as the 21-22Us described below were only found in the 5’pN enriched library but not 5’ modification-independent library, highlighting the importance of using multiple library preparation methods to investigate sRNA expression.

### 21-22U RNAs with a 5’ monophosphate resembling piRNA target TEs associated with the parasitic life cycle stage

After miRNAs, sRNAs originating from intergenic spaces, protein-coding genes and TEs were most highly expressed group of sRNAs in the 5’pN-enriched library sequences (**Figure 1a**). We further investigated these sRNAs to identify specific classes of sRNAs upregulated in the PF and FLF. Analysis of the length and first nucleotide at the 5’ site revealed that 21-22 nt long sRNAs starting with uracil (hereon in referred to as 21-22Us) were the most highly expressed 5’pN sRNA upregulated in PF (**Figure 2a**). In contrast, sRNA sequences with a 5’ pN either upregulated in the FLF or not differentially expressed (DE) showed no bias for a particular length or propensity for a particular 5’ base (**Figure 2a**). In total, we identified 1887 unique 21-22U sequences upregulated in the PF, 86 sequences in the FLF and 218 sequences non-differentially expressed (non-DE), respectively. Given the larger number of unique 21-22U sequences and the higher expression levels in the PF, we propose that the 21-22Us are a class of sRNAs with a role in parasitism or a feature associated with the parasitic life cycle stage.

**Figure 2:**
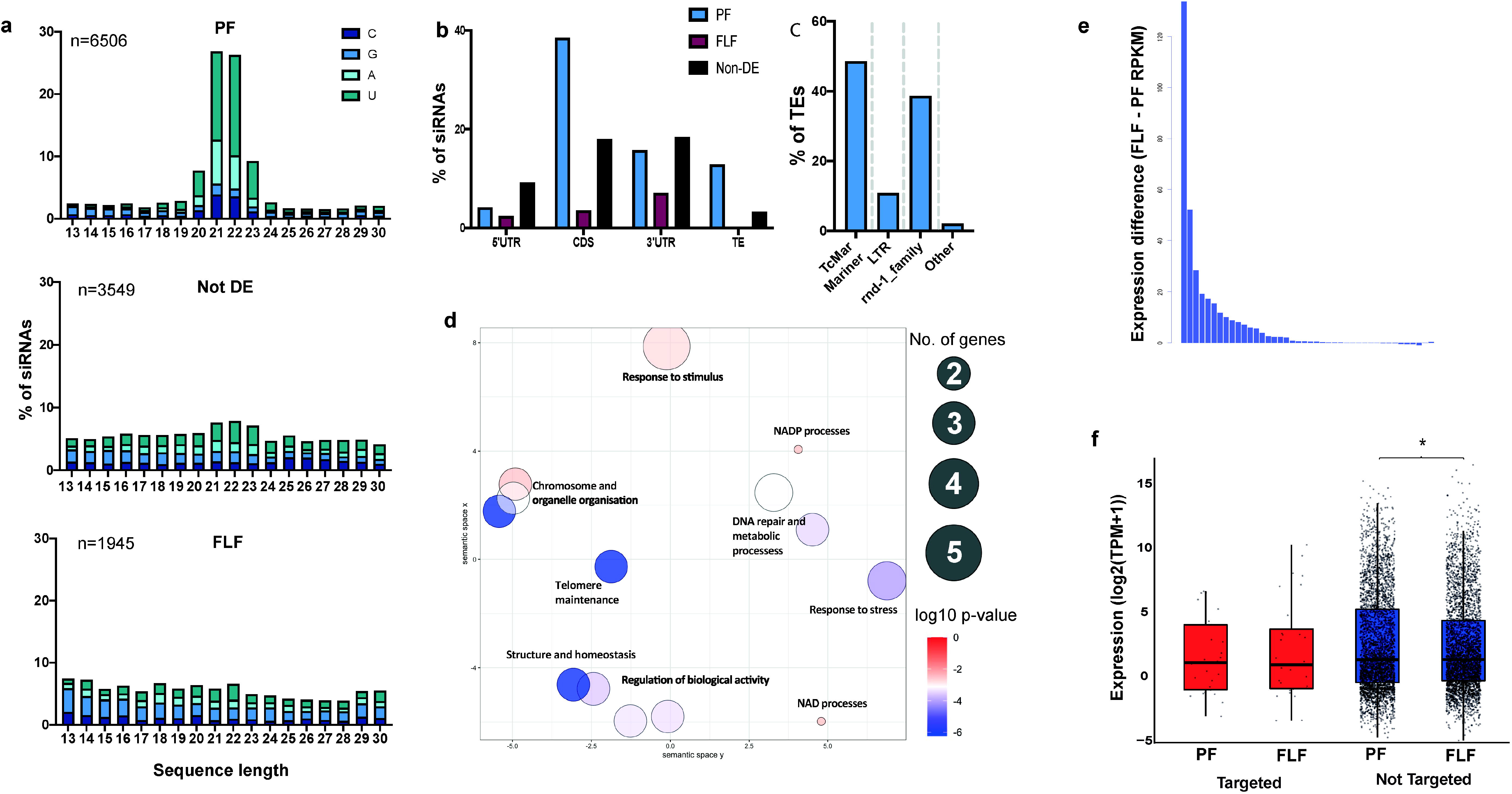
21-22Us with a 5’ monophosphate modification associated with parasitism. (a) Length distribution and 5’ starting nucleotide of 13 – 30 nt sRNAs originating from either protein-coding genes, intergenic spaces or transposable elements with a 5’ monophosphate modification, upregulated in the PF (n=6506 sequences), FLF (n=1945 sequences) or not DE (n=3549 sequences). (b) Predicted targets of significantly upregulated 21-22Us in the PF, FLF and non-DE based on antisense sequence complementarity. Targets include protein-coding regions (CDS, 5’UTR and 3’UTR) and TE sequences. (c) Classification of TE targets of 21-22Us in the PF. Targets include TcMar-Mariner DNA transposon (48.5%), LTR retrotransposons (10.8%) and an unclassified family (38.6%). (d) Enriched GO terms based on the biological function (BF) of the 21-22U targeted gene. Clustering of the GO terms based on similar functions was carried out using REVIGO. Size of circle indicates the number of genes and colour indicates p-value. (e) Difference in expression level (RPKM: FLF minus PF) of genes targeted by PF-upregulated 21-22Us. The targeted genes were more highly expressed in the FLF *cf.* PF life cycle stage. (f) Expression of TEs either targeted or not targeted by PF-upregulated 21-22Us. * indicates p<0.01.

### The 21-22Us target TE-associated protein coding-genes and TE sequences

Based on sequence complementarity, we predicted the targets of 21-22U RNAs upregulated in the PF. Of the 1887 unique 21-22U sequences, 726 targeted the coding sequence (38.47%), followed by the 3’ UTR (15.69%) and 5’ UTR (4.13%) regions of 42 *S. ratti* protein-coding genes (**Figure 2b**). Of those 42 protein-coding genes, 13 (30.9%) had predicted functions (**Supplementary Data 4)**, that were directly associated with TE activity including DNA helicase, helitron-like proteins, reverse transcriptase and transposase-encoding genes. In addition, 12 target genes (28.6%) were classed as ‘hypothetical’ and were not annotated with a function (**Supplementary Data 4**). To characterise the function of the hypothetical proteins, we grouped them into orthofamiles using Orthofinder2 ^29^ with 18 other nematode species **(Supplementary Data 5)**. Interestingly, we found that ten of the twelve hypothetical genes were *S. ratti*-specific. Only two genes were found in other species, one which belongs to the Mos1 transposases family and the second which did not have a known function in related species. The target genes were significantly enriched (Fishers Exact Test FDR < 0.01) for Gene Ontology (GO) terms associated with chromosome and telomere organisation and maintenance, cellular responses to stress and DNA damage and homeostatic processes (**Figure 2c, Supplementary Data 6**). We found that the genes targeted by 21-22Us were expressed at higher levels in the FLF compared with PF suggesting that 21-22Us may repress gene expression (**Figure 2d**). In addition to the protein-coding genes, ten miRNA precursor sequences and two tRNA genes were also perfectly antisense, and therefore likely to be targeted by PF-upregulated 21-22U RNAs (**Supplementary Data 4**).

Because many of the protein-coding gene targets were associated with TE-activity, we sought to investigate if 21-22Us are involved in directly targeting and regulating the expression of TE sequences. We first improved the annotation of TE sequences within the *S. ratti* genome (See Methods, **Supplementary Figs. 2-4, Supplementary Data 7**) and identified TE sequences that are perfectly antisense to 21-22Us. Our results showed that 12.8% of PF-upregulated 21-22U RNAs targeted the DNA transposon TcMar-Mariner (48.5% of the TEs targeted), followed by an unannotated class of TEs (38.6% of the TEs targeted) and long terminal repeats (LTR) retrotransposons (10.8% of the TEs targeted) (**Figure 2e, Supplementary Data 8**). These results suggest that the main role of the PF-upregulated 21-22Us is the regulation of TE activity (**Table 1**), mainly of the DNA transposon family. We then examined the expression of 21-22U-targeted TE transcripts and non-targeted TEs in the PF and the FLF. Overall, we found that the expression level of TEs targeted by 21-22Us was similar in the PF and FLF (**Figure 2f, Supplementary Data 9**).

### 21-22Us and their TE targets are physically clustered on the X-chromosome

To examine the distribution of the 21-22Us upregulated in the PF, we mapped the sequences to the *S. ratti* genome. The 21-22U loci clustered on the second largest scaffold of the X-chromosome, forming two main clusters across both strands of the genome spanning 1 and 2.7Mb, respectively (**Figure 3a, Table 1**). In total, 53.27% of the PF-upregulated 21-22Us mapped to this X-chromosome scaffold. For comparison, we also mapped the comparatively small number of 21-22Us sequences upregulated in the FLF and non-DE datasets to the genome, and these showed no evidence of large clustering (**Supplementary Fig. 5)**. The TEs targeted by 21-22Us in the PF were also predominantly located on the X-chromosome, but from different regions (**Figure 3b**). This is in contrast to the distribution of TEs in general which are found throughout the genome (**Supplementary Fig. 6**). Further investigation of 21-22U loci in the genome revealed the sequences originate from overlapping same-strand clusters (found across both strands) in the genome (**Figure 3c**). In total, 88.78% of 21-22Us loci overlapped with at least one other 21-22U sequence on the same strand of the genome, and on average each 21-22U sequence overlapped with 13.9 ± 0.28 (mean ± SE) sequences (**Figure 3c**), which is similar to the pattern observed for piRNAs in *C. elegans* ^30^.

**Figure 3:**
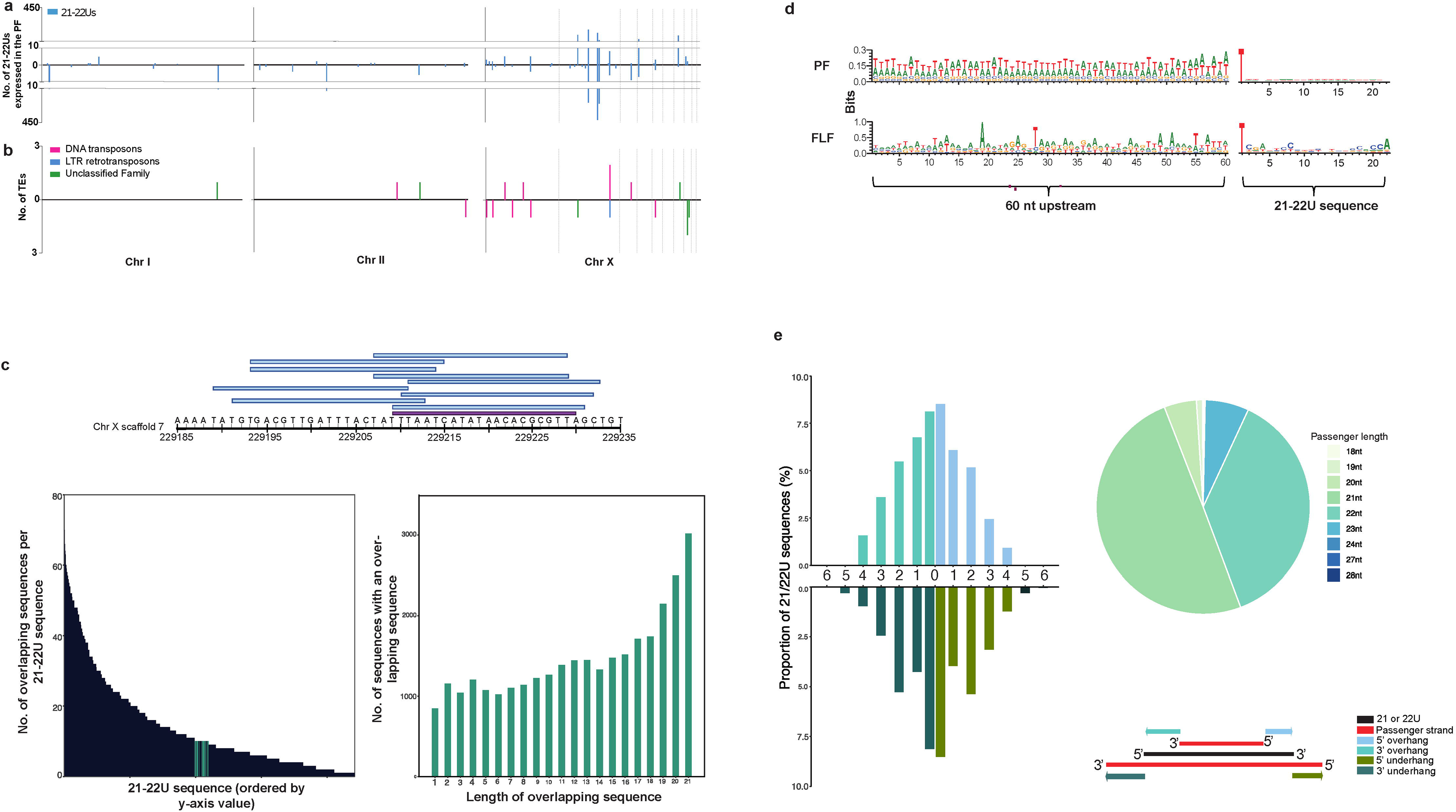
Chromosomal distribution and overlapping patterns of 21-22Us which is similar to *C. elegans* piRNAs. (a) Distribution of upregulated 21-22Us across the genome which consists of two autosomes (11.7Mb chromosomes I and 16.7Mb chromosome II) and the chromosome X (12.9Mb made up of 10 scaffolds). (b) Distribution of 21-22U TE targets across the genome (n = 20 TE targets). (c) Identification of an overlap signature in the 21-22Us. Figure showing an example with a 21Us originating from chromosome X and all overlapping 21-22U strands. Bottom left figure showing the number of 21-22Us that overlap with themselves (88.78%). Bottom right figure showing the overlap lengths of siRNAs against the 21-22Us. (d) Sequence logos of PF 21-22U upstream sequences versus the FLF 21-22Us to identify nucleotide richness based on the bits of each nucleotide. (e) No dicer-processing signature in PF 21-22Us, based on the overhang and underhang of the passenger strand. Passenger strands cleaved by the enzyme Dicer, leave a distinguishable 2-3’ overhang. Pie chart shows the percentage of passenger strands

### Upregulated 21-22Us in the PF have AU rich upstream sequences

We have determined that the 21-22Us expressed by *S. ratti* PF share similar features with *C. elegans* piRNAs including a similar length and first 5’ nucleotide, targeting of TEs and they originate from overlapping sequences in the genome (**Table 1**). We further investigated *S. ratti* 21-22Us for features that are associated with *C. elegans* piRNAs, namely, an AT rich upstream sequence and a conserved CTGTTTCA motif upstream of the 21U loci found in type I piRNAs ^30 31^. We identified AT richness in the sequence upstream of *S. ratti* 21-22Us comparable to *C. elegans* piRNAs (**Figure 3d**). However, a piRNA-associated motif was not found upstream of the *S. ratti* 21-22U loci.

### 21-22Us are not Dicer-processed

Classes of sRNA can be characterised by the mechanism used to process mature sRNA sequences from double stranded RNA precursor sequences. We searched for Dicer-processing signatures in 21-22U sequences, which can be identified by a 3’overhang in sRNA duplexes. However, a Dicer-signature was not observed for 21-22Us and the profile of sRNA duplexes more closely resembled patterns observed for RdRP-processing (**Figure 3e**). Furthermore, no evidence of a ping-pong signature was observed for 21-22Us (**Table 1**), which is usually associated with piRNAs in *D*. *melanogaster* showing a 10 nt overlap of the 5′ ends of the piRNAs with other sRNA sequences (**Supplementary Fig. 7**).

### Distinct subsets of 27GAs with a 5’ polyphosphate are predicted to target TEs in the PF and FLF

We sought to identify 5’-triphosphated sRNAs (potential RdRP siRNAs) in *S. ratti* by removing the sequences that were present in the 5’pN-enriched libraries from those in the 5’ modification-independent libraries. Expression analysis after the subtraction revealed that the most highly expressed sRNAs in the PF were 27 nt long (hereinafter referred to as 27GAs) (**Figure 4a**). A subset of 27GAs was also significantly upregulated in the FLF (**Figure 4a**). In total, we identified 14,292 unique 27GAs including 193 upregulated in PF and 52 in FLF. Expression of 27GAs was previously reported for mixed-sex free-living adults ^7^, but here we found that 27GAs are also expressed in the PF, and that distinct subsets of 27GAs are associated with PF and FLF life cycle stages.

**Figure 4:**
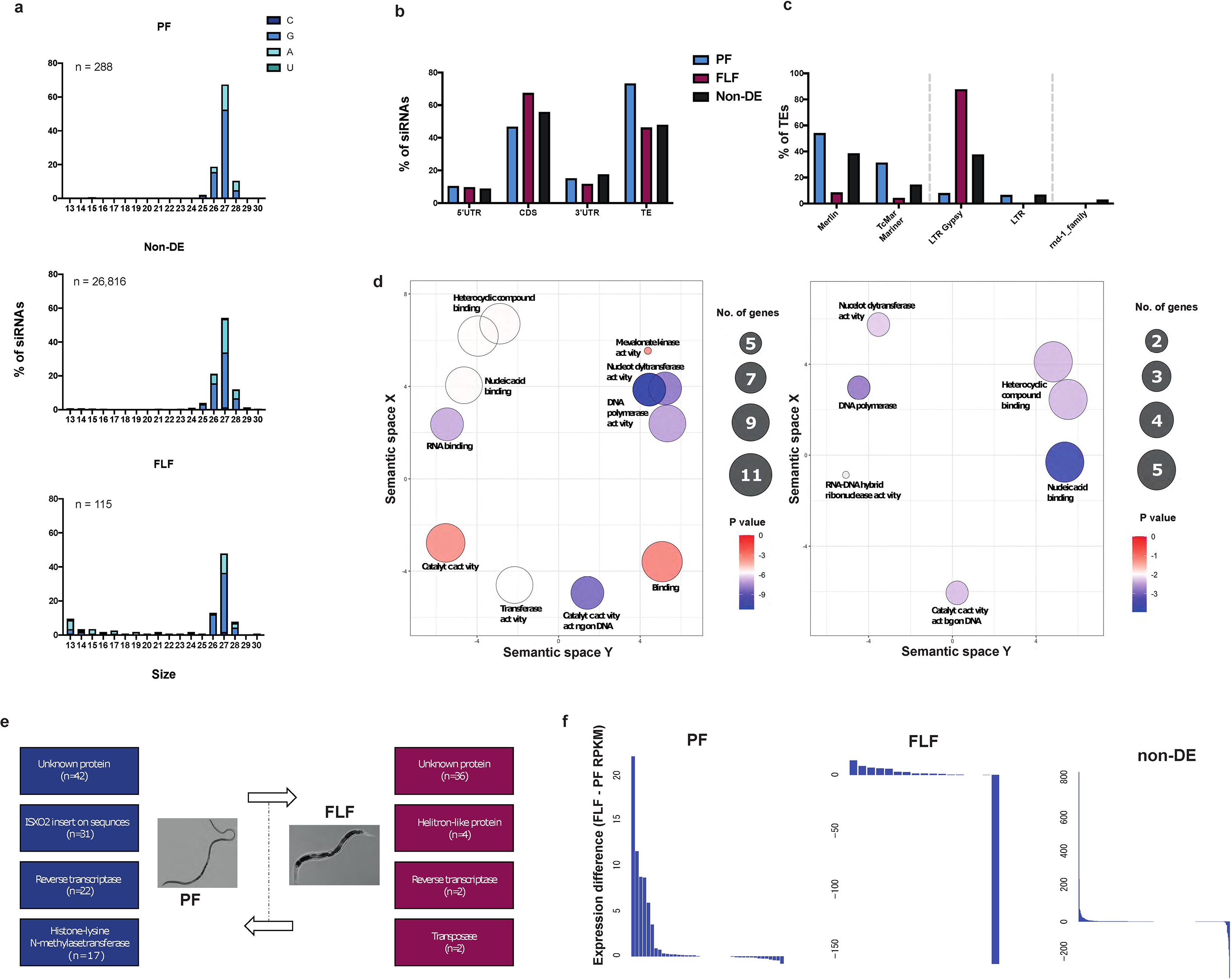
27GA 5’-polyphosphate sRNAs upregulated at different life cycle stages. To identify sRNAs with a 5’ modification, sRNA sequences found in a 5pN library were filtered out from sRNAs identified in the 5’ modification-independent library. (a) Length distribution and 5’ starting nucleotide of 13 – 30 nt sRNAs with a 5’ modification, upregulated in the PF (n=288 sequences), FLF (n=115 sequences) or those not DE (n=26,816), excluding miRNAs, tRFs and rsRNA sequences. 27GAs are most abundant among them. (b) Putative targets of the 27GAs in the PF, FLF and non-DE. Target protein-coding regions (CDS, 5’UTR and 3’UTR), TE were predicted based on antisense sequence complementarity. (c) Classes of TE antisense to PF-upregulated, FLF-upregulated and non-DE 27GAs. (d) Enriched GO terms of the genes targeted by PF-upregulated 27GAs (left) or FLF-upregulated 27GAs (right). Clustering of each GO term (biological function) based on similar functions was carried out using REVIGO. Size of circle indicates the number of genes and colour indicates p-value. (e) Predicted protein function of target genes and the number (n) of 27GAs that target. (f) Difference in expression level (RPKM: FLF minus PF) of genes targeted by PF-upregulate, FLF-upregulated and non-DE 27GAs

### 27GA RNAs target TE-associated genes

The target sequences of 27GAs were predicted based on antisense. Of the 193 and 52 27GAs upregulated in the PF and FLF, the majority of 27GAs targeted the coding sequence (46.63% and 67.31%) followed by the 3’UTR (15.03% and 11.54%) and 5’UTR (10.36% and 9.62%) of protein-coding genes **(Figure 4b)**. A similar pattern was observed for the 14,047 non-DE 27GAs, which were targeting the coding sequence (55.64%), followed by the 3’UTR (17.41%) and the 5’UTR (8.73%) **(Figure 4b)**. In total, the PF-upregulated 27GAs targeted 40 protein-coding genes, FLF-upregulated 27GAs targeted 17 protein-coding genes and non-DE 27GAs targeted 452 protein-coding genes. No potential targets were identified for rRNA, tRNA and miRNA precursors sequences. The target genes across all three subsets, PF-upregulated, FLF-upregulated and non-DE, were predominantly associated with TE activity including transposases, reverse transcriptase, helicase, helitron integrase-coding genes **(Supplementary Data 10)**. The PF-upregulated, FLF-upregulated and non-DE gene targets were enriched for similar Biological Processes (BP) and Molecular Function (MF) GO terms (**Supplementary Data 11**) including GO terms associated with DNA integration and biosynthesis, RNA binding and DNA polymerase activity (**Figure 4c, Supplementary Data 11**).

However, the specific genes targeted by either the PF-upregulated or the FLF-upregulated 27GAs were different. PF-upregulated 27GAs targeted genes coding for transposase insertion sequence XO2 (31 sRNAs targeted 11 genes, compared to only one gene targeted by FLF-upregulated 27GAs) known to mediate transposition, and reverse transcriptase related genes (22 sRNAs targeted 7 genes) (**Figure 4d**). A large proportion of genes targeted by the PF-upregulated and FLF-upregulated 27GAs coded for ‘hypothetical’ proteins (PF: 42 27GAs targeting 13 genes, FLF: 36 27GAs targeting 7 genes; **Supplementary Data 10)**. The PF-upregulated 27GAs are targeting genes belonging to two orthofamilies. Eight genes belong to an orthofamily comprising genes encoding ISXO2 transposase family and three genes belong to an orthofamliy comprising Mos1 transposase genes, both of which are related to TE activity. The FL-upregulated 27GAs targeted one gene belonging to an orthofamily coding for genes related to nucleic acid binding activity (**Supplementary Data 5**). Together, these results support the role of 27GAs in TE regulation, and that specific subclasses of TEs are targeted at different life cycle stages. A notable difference between the sets of target genes is that the PF-upregulated, but not FLF-upregulated 27GAs, targeted four histone-lysine N methyltransferase-coding genes, all located on the X-chromosome. Expression level of genes targeted by PF-upregulated 27GAs were lower in the PF compared with FLF. The non-DE 27GA-targeted genes were expressed at similar levels in the PF and FLF (**Figure 4e**).

### 27GAs target TE sequences

The 27GAs previously identified in the FLF were predicted to target TE sequences ^7^. Our analysis above further shows that 27GAs expressed in the PF and FLF are targeting protein-coding genes associated with TE activity. Here, we also investigated if PF-expressed 27GAs directly target TE sequences (**Figure 4b, Supplementary Data 12**). We identified 27GAs that aligned perfectly antisense to class I and class II TEs in the *S. ratti* genome and found that a distinct set of TEs were targeted by the PF and FLF. Our results showed that 73.06% (n= 141) of PF-upregulated 27GAs were predicted to target 42 TE sequences, many of which were DNA transposons. Of the PF-upregulated 27GAs targeting TEs, 53.9% were antisense to the DNA transposon from the Merlin family and a further 31.2% targeted TcMar-Mariner. In addition, 7.8% and 6.4% targeted the retrotransposons LTR gypsy family and LTRs of unannotated families, respectively (**Figure 4f**). In comparison, 46.15% (n= 24) of the FL-upregulated 27GAs and 47.83% (n= 6719) of the non-DE 27GAs also targeted TEs, but the overall proportion of these sequences that targeted TEs was lower compared to PF-upregulated TEs. Unlike the 27GAs upregulated in the PF, FLF-upregulated 27GAs predominantly targeted the class I retrotransposons LTR gypsy (87.5% of the TEs targeted), followed by the DNA transposons Merlin (8.3% of the TEs targeted) and TcMar-Mariner (4.2% of the TEs targeted). In the non-DE 27GAs several TE families were targeted including Merlin (38.4%), LTR gypsy (37.5%), TcMar-Mariner (14.4%), LTRs belonging to an unannotated subfamily (6.6%), unclassified TE families (2.8%) and LTR copia (0.08%) (**Figure 4f**). Together, these results indicate that 27GAs in the PF are important in targeting, and presumably regulating, the activity of TEs within the DNA transposon family, in comparison to the FL-upregulated and non-DE 27GAs which also target and regulate retrotransposons (**Table 1**). To investigate the role of TEs further, we compared the expression of TE sequences that are targeted vs. not targeted by the different subsets of 27GAs (PF-upregulated, FLF-upregulated and non-DE) (**Supplementary Fig. 9**). We found that the TE sequences targeted by either PF-upregulated or FLF-upregulated 27GAs were expressed at similar levels to non-targeted TEs (**Supplementary Data 9**). The TEs targeted by non-DE 27GAs were expressed at lower levels compared to the non-targeted TEs. A significant difference was found in the TE expression level between PF and FLF for targeted TEs. However, the difference in expression level was not clearly directional and included TEs that were expressed at either higher or lower levels in the PF (**Supplementary Fig. 9**), demonstrating the diversity in TE expression levels, and presumably their regulation, across the two life cycle stages.

### 27GAs are clustered within the X-chromosome

We investigated the genomic distribution of 27GAs by mapping the 27GAs sequences to the *S. ratti* genome. Similar to the 21-22Us, the 27GAs were predominantly located on the X-chromosome (**Figure 5**). However, unlike 21-22U RNAs that were clustered in one particular region of the X-chromosome (**Figure 2d**), the distribution of the 27GAs spanned across most of the X-chromosome scaffolds (**Figure 5**). Of the PF-upregulated 27GAs, 21.24% were located on the 3^rd^ largest X-chromosome scaffold, followed by 12.95% on the second largest X-chromosome. In comparison, the largest cluster of 15.38% of FLF-upregulated 27GAs were located on the opposite strand of the 6^th^ largest scaffold of the X chromosome spanning the first 100 Kbp. The non-DE 27GAs were found throughout the X-chromosome as well as chromosome I and II (**Figure 5**). These results show that although TEs in general are distributed across the *S. ratti* genome (**Supplementary Fig. 6**), the TEs targeted by 21-22Us and 27GAs were predominantly located on the X-chromosome, indicating that *S. ratti* sRNAs are mainly involved in regulating X-chromosome-specific TEs (**Supplementary Fig. 6**, **Supplementary Fig. 9**, **Figure 3**).

**Figure 5:**
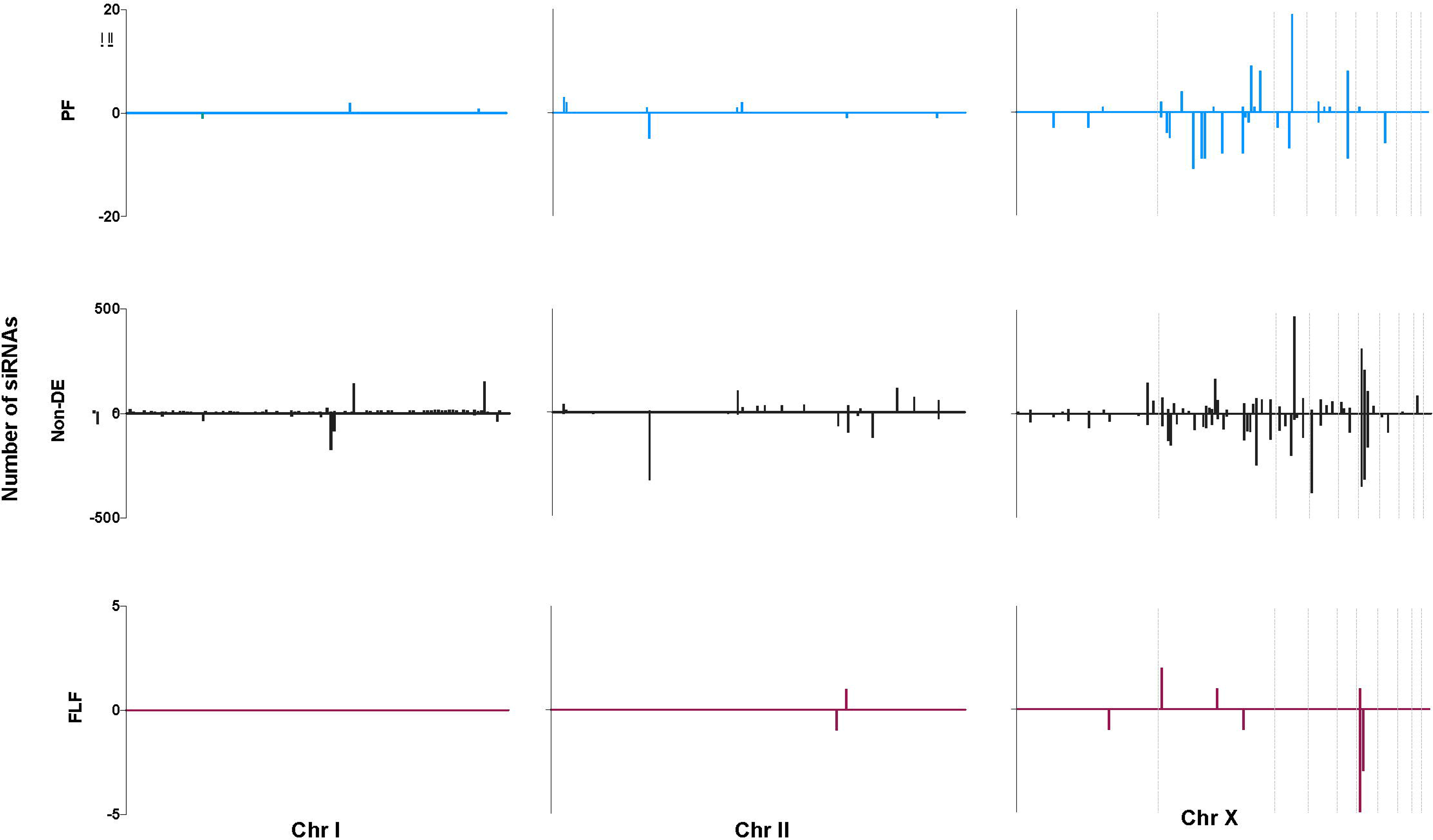
Chromosomal distribution of upregulated 27GAs in the PF (n=193 sequences), FLF (n=52 sequences) and non-DE (n=14,047 sequences) within I (11.7Mb), II (16.7Mb) and the X chromosome (13Mb), identified using sequence complementarity

### Subsets of miRNAs are differentially expressed by PF and FLF life cycle stages

Using Mirdeep2 ^32^, we identified a total of 158 miRNAs, including 103 and 94 miRNAs expressed across both replicates of either the PF or FLF life cycle stage, respectively. The majority of miRNAs we identified had previously been reported in miRBase v22 ^33^, however, we identified four novel miRNAs (**Supplementary Data 13**). To investigate which miRNAs are most likely to have a role in parasitism, we identified miRNA sequences that were upregulated in the PF or FLF life cycle stages. Nine and six miRNA sequences were significantly upregulated in the PF and FLF life cycle stages, respectively (edgeR FDR< 0.01, **Figure 6a, Supplementary Data 13**).

**Figure 6:**
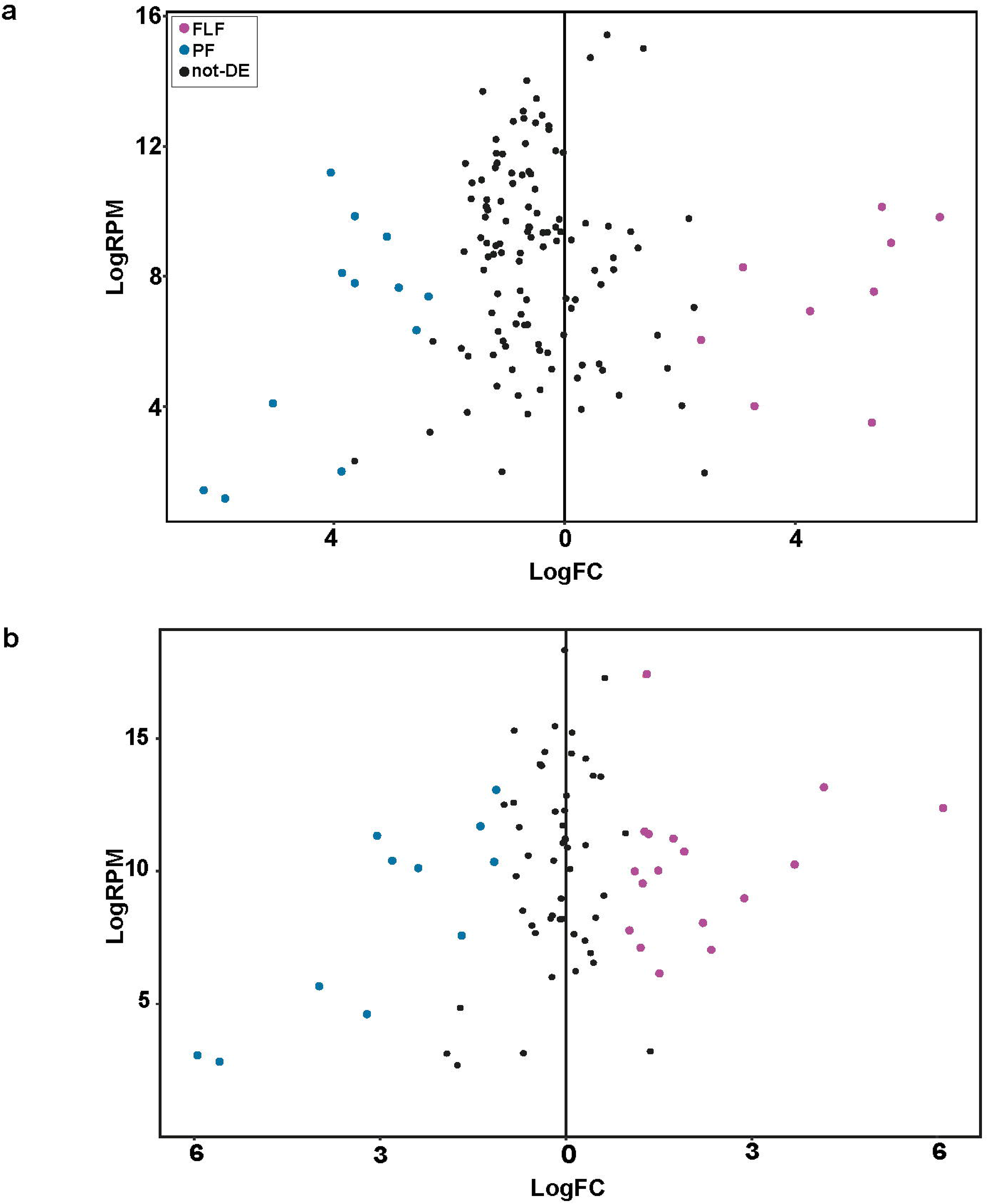
Differential expression of (a) miRNA sequences showing miRNAs upregulated in the PF (n=12 sequences), FLF (n=9 sequences) (FDR< 0.01) and (b) seed sequences

We categorised the *S. ratti* miRNAs into families based on their seed sequences and identified a total of 92 seed families (**Supplementary Data 14**). Comparison of the expression levels based on seed revealed that 23 and 17 seed sequences were significantly upregulated in PF and FLF, respectively (**Figure 6b, Supplementary Data 14**). The seed families that were differentially expressed comprised between 1 and 21 miRNA sequences and included both miRNA families conserved across other species and uncharacterised seed families. The miRNAs in the seed family with the most members (seed sequence UUGCGAC) were predominantly upregulated in the PF and may therefore target a specific set of mRNAs important in parasitism. The UUGCGAC miRNA family was not found in seven other nematode species where data was available on miRBase (*Ascaris suum, Brugia malayi, C. elegans, Haemonchus contortus, Heligmosomoides polygyrus, Pristionchus pacificus* and *Panagrellus redivivus*) indicating that it is likely to be a *Strongyloides*-specific family.

## Discussion

We have investigated sRNAs in the gastrointestinal parasite *S. ratti* by directly comparing sRNAs expressed in genetically identical PF and FLF life cycle stages. We have identified two distinct classes of sRNAs, characteristic of siRNAs, predicted to target TEs including a (i) piRNA-like 21-22Us with a 5’pN, highly associated with the PF life cycle stage, and (ii) 27GAs with a 5’ modification, which have distinct subsets of sequences upregulated in the PF and FLF. We have also identified miRNA sequences and miRNA families based on their seed sequence, that are differentially expressed in the PF and FLF. We propose that sRNAs expressed at higher levels in the PF are either directly related to parasitism or related to a feature associated with the parasitic generation.

### Parasitism-associated 21-22U sRNAs target TEs and resemble piRNAs

We identified 1887 unique 21-22Us significantly upregulated in the PF. We postulate that these 21-22Us are particularly related to the *S. ratti* adult parasitic stage of the life cycle because high expression levels of this class of sRNAs were not observed in FLFs in this study or in other life cycle stages previously investigated ^7^. We predicted the sequences that were targeted by 21-22Us based on perfect antisense complementarity. Collectively, our results strongly support that 21-22Us are targeting TEs. sRNAs that regulate TEs are usually most highly expressed in the germline in *C. elegans* and other animals ^1,49^, and it is therefore likely that the TEs and 21-22Us expressed here are from the PF germline cells. Interestingly, the TEs targeted by PF-upregulated 21-22Us are expressed at similar levels in the PF and FLF and compared to TEs not targeted by 21-22Us. If we assume, based on evidence in other animals, that the expression of TEs targeted by sRNAs are repressed ^34^, then it is likely that 21-22Us could be acting to repress the expression of a subset of highly expressed TEs back to the ‘normal’ levels observed in FLF and non-targeted TEs. It is important to note here that the analysis of TE transcript activity was based on polyA-selected RNAseq data and therefore is only informative about polyadenylated TEs e.g. retrotransposons with a polyA sequence and the transposase component of DNA transposons.

In *C. elegans, D. melanogaster* and *M*. *musculus,* piRNAs have a key role in regulating and silencing TE activity ^35,67^. Given the similarity in size (21-22 nt), 5’ nt (uracil), 5’ monophosphate modification, no Dicer-processing signature and their predicted targeting of TEs, we investigated if other features associated with piRNAs were also common to *S. ratti* 21-22Us. The piRNAs found in *C. elegans* originate from large genomic clusters that give rise to short 21U piRNAs. In *C. elegans,* piRNAs can be further divided into two groups: type I 21U RNAs that make up 95% of total piRNAs and the less abundant type II 21U RNAs ^31,35,36^. Type I piRNAs are transcribed from thousands of AT rich loci that accumulate within two large clusters on chromosome IV and have a conserved upstream ‘CTGTTTCA’ Ruby motif ^31,37^. These clusters mainly originate from introns and intergenic regions that overlap with other 21Us ^3031^. In contrast, Type II 21Us are distributed throughout the genome and have no upstream motif ^30,36^. *S. ratti* 21-22Us resemble *C. elegans* type I piRNAs, because they (i) are clustered in a specific region of the genome, though they are located on the X-chromosome rather than an autosome, (ii) have an AT rich upstream sequence, (iii) share a similar, same-strand overlapping pattern in the genome for the 21-22U loci. Like *C. elegans* piRNAs, the number of nucleotides overlapped are varied, but most commonly, 21-22Us overlapped by 20-21 nucleotides with other 21-22Us. Interestingly, the only validated target of the *C. elegans* piRNA pathway is the DNA transposon family, Tc3^13^. We have shown that *S. ratti* 21-22Us also predominantly target DNA transposons. Together our results indicate that *S. ratti* 21-22Us may be the equivalent to the *C. elegans* piRNAs and processed through a similar manner.

An intriguing difference between *S. ratti* 21-22Us and *C. elegans* 21Us, is that *S. ratti* 21-22Us share characteristics associated with piRNAs in *D. melanogaster* not found in *C. elegans*. *D. melanogaster* piRNAs are transcribed from piRNA clusters made up of transposons or repeat elements and are characterised as repeat associated siRNAs (rasiRNAs) ^15,38,39^. These rasiRNAs highly target transposons through complementary base pairing and are required for the normal development of the germline ^16,40^. A similar pattern was observed in *S. ratti* 21-22Us. Although we have shown that 21-22Us originate from clusters found on the X-chromosome, some of these clusters are found in intergenic regions, however, like *D. melanogaster,* 21-22Us also originate from TEs, as well as predominately targeting TEs clustered on the X-chromosome through perfect antisense complementarity. This indicates that 21-22Us may be derived from TE regions that are required to be regulated and silenced, similar to the piRNAs of *D. melanogaster.*

Collectively, our results suggest that *S. ratti* 21-22Us share many similarities with the piRNA class of sRNAs, a pathway which is assumed to have been lost in nematodes outside of clade V ^5^. It is possible that the 21-22Us may have originated from an ancestral PIWI pathway which has now lost key components such as the PIWI Argonautes, and the pathway has subsequently diverged in *S. ratti*. However, the lack of an upstream Ruby motif that is found for *C. elegans* piRNAs or the ping pong signature in *D. melanogaster* suggests that this is either not the case or the 21-22Us derived from piRNA substantially long enough time ago. Another possibility is that many of the features common to piRNA and 21-22Us are universally advantageous to TE targeting by sRNAs and they have evolved independently. Furthermore, a lack of PIWI Argonaute-coding genes in *S. ratti* suggests that an alternative pathway is associated with 21-22Us. Indeed, an alternative class of sRNA for silencing TEs has been proposed for other nematodes lacking the PIWI pathway ^5^. However, studies on the alternative classes of TE-targeting sRNAs in nematodes have not reported similarities with piRNAs, as is the case for the 21-22Us in this study. Previously, we have identified a group of Argonaute-coding genes closely related to *C. elegans* WAGOs that are significantly upregulated in the PF compared with the FLF life cycle stages in four *Strongyloides* species investigated ^6^, and we speculate that based on their expression patterns these could be associated with 21-22Us in *S. ratti*. Further work is required to identify if these Argonaute proteins are associated with 21-22Us and to identify other components of this pathway.

### 27GAs are predicted to regulate TEs in both the parasitic and free-living adult stages

We identified a second class of sRNAs, the 27GAs, that also target and putatively regulate the expression of TEs. In contrast to the 21-22Us, 27GAs were expressed across both life cycle stages, and were only observed in the 5’ modification-independent library, suggesting that they possess a polyphosphate or capped 5’ modification. The 27GAs have previously been described in adult free-living *S. ratti* worms (mixed female and male) ^7^ but this is the first time they have been identified in the PF. In addition to the 27GAs that were expressed at similar levels in both life cycles stages, smaller subsets of 27GAs were upregulated in the PF or FLF. Although 27GAs in both the PF and FLF were targeting TEs, there was a clear difference in the classes of TEs targeted. The PF-

upregulated 27GAs mainly targeted TEs from the class II DNA transposons, namely Merlin and TcMar-Mariner, whereas the FLF-upregulated 27GAs predominantly targeted the class I retrotransposon LTR gypsy. However, it is not clear why there is a difference between the type of TEs targeted in the PF and FLF. Protein-coding genes targeted by the 27GAs were associated predominantly with TE activity but some are with other biological processes. For example, the PF-upregulated 27GAs target histone-lysine-n-methyltransferase-coding genes. These proteins are involved in histone modification and play a role in chromatin structure and gene expression ^41^. Inhibiting histone methyltransferase has been shown to stop the life cycle of *Schistosoma mansoni*, suggesting that the role of histone methylation is extremely important in parasites where differentiation of the life cycle occurs within a host ^41^. There could therefore be a specific role of 27GAs in regulating histone modification that is specific to the PF life cycle stage.

Unlike the 21-22Us, the 27GAs share few similarities with piRNAs. Instead the 27GAs are more similar to the secondary 22G siRNAs in *C. elegans* which have a 5’ triphosphate, a 5’ guanine (5’G) bias and are processed by RdRP ^42,43^. The 27GAs, showed no evidence for Dicer-processing further supporting that they are likely to be processed in a similar manner to *C. elegans* 22Gs RdRPs. sRNAs similar to secondary 22Gs in *C. elegans* have also been reported in other nematodes. For example, the clade I-III nematodes produce 22G sRNAs processed by RdRPs, which mainly target TEs in the absence of piRNAs ^5,44^. RdRP orthologues related to the processing of 22G siRNAs in *C. elegans* are also present in *Strongyloides* ^7^, indicating that the RdRP pathway is active.

### Why is TE regulation important in the parasitic stage?

Overall, our results have shown that expression of sRNAs that target TEs in *S. ratti* are expressed at higher levels in the PF *cf.* FLF, including the 21-22Us and 27GAs. This suggests that the regulation of TE activity is higher in the PF, raising the question, *why is the regulation of TE higher in the PF stage?* It is widely accepted that TE activity is often associated with the germline ^45,46^. Most TE insertions are considered to be detrimental to the genome integrity and leading to disruption of genes or regulatory regions ^11^. However, TE activity may also be beneficial and can be related to genome rearrangement, regulation of genes and chromosome stability ^11,47^. For example, LINE-like retrotransposons are highly important and required for the maintenance of telomeres in *Drosophila,* where the telomerase enzyme is missing ^48^. From our study, it is unclear if the higher level of TE regulation observed in PF is beneficial to the *S. ratti* genome and this requires further investigation. The increase in TE regulation in the PF could reflect the stressful environment that *S. ratti* is exposed to in the rat host, and the increased sRNA activity is a direct response to regulate the activity of increased TE activity. It is also possible that differences in expression level between PF and FLF represent differences in the proportion of germline cells in each life cycle stage. For example, if PFs have a larger proportion of germline cells compared with FLF, this could be observed as higher level of expression of germline-related transcripts e.g. TEs or sRNAs.

In addition to differences in lifestyle i.e. parasitic stages inside the host vs. free-living stages outside of the host, PF and FLF also differ in their reproductive strategy; the PF reproduces through mitotic parthenogenesis and the FLF reproduces through sexual reproduction ^49^. The differences observed between TE activity and the subsequent TE regulation by sRNAs could reflect differences in embryogenesis and development. TEs have also been associated with a role in cis regulation of genes involved in sexual development ^50^. The 21-22Us, 27GAs and the TEs that they targeted are predominantly located on the X-chromosome (this is not the case for TEs in general). TEs are often associated with sex chromosomes and in some cases these TEs have a role in regulating sex-chromosomal genes ^50^. With the results presented in this study and the presence of two genetically identical life cycle stages with different reproductive strategies, *S. ratti* offers a particularly interesting study platform to investigate the role of TEs and sRNAs in reproduction, sexual development and the regulation of sex-chromosomal genes. Lastly, TEs are fast evolving sequences subject to higher mutation rates compared with other sequences in the genome. It is therefore important that a regulatory system can respond and adapt rapidly to variation in these sequences to accurately regulate active TEs. In this study we have looked at a single time point in the PF and FLF life cycles stages. The 27GAs upregulated in either the PF or FLF represent a response to the TEs that are upregulated in that snapshot of time. However, the TEs that are active in the genome at any one time may vary and the 27GAs would subsequently vary in response to these TE sequences. Interestingly there are clearly distinct trends between the TEs and TE-associated genes targeted by the PF and FLF and this could mean that particular classes or TE are more likely to be active in either life cycle stage.

### Other classes of sRNA are differentially expressed

Our results identified a distinct set of miRNAs and miRNA families upregulated in the PF and FLF. Unlike siRNAs which require perfect or near-perfect complementarity to their target, miRNAs have just a small seed sequence of ~7 nucleotides and bioinformatic prediction of miRNA targets is thus prone to false positives and was therefore not addressed here. We have previously identified protein-coding genes upregulated in the PF *cf.* FLF stage that have a putative role in parasitism including genes that are physically clustered in the genome ^2,76,51^. The differentially expressed miRNAs may be involved in regulating these ‘parasitism genes’ and further lab-based approaches are required to investigate this.

We also observed a difference in the expression levels of sRNAs originating from mature tRNAs known as tRNA derived sRNA fragments (tRF) ^52,53^. Mature tRNAs can be cleaved to produce several classes of tRF classified as 5’ and 3’ tRFs, 5’ and 3’ tR-halves, 3’ CCA-tRF and internal-tRF. We analysed tRFs found in the PF and FLF, and found that there is an increased expression level of internal-tRFs in the FLF. The tRFs have important biological roles in the translation of genes as well as gene regulation through the interaction with proteins and mRNAs ^54^. Their expression has been shown to be upregulated during stress and starvation in *Trypanosoma brucei* ^55^, which suggests that increased expression in the FLF may be due to the stressful conditions of the environment. tRFs have not been well characterised and relatively little is known about this class of sRNA. More work in this area is required to elucidate the role of tRFs. The library preparation methods used here enrich for RNA molecules with a 5’ monophosphate which are likely to represent ‘true’ sRNAs, however, we cannot rule out that some of the reads represent degraded products of tRNA or other longer RNA molecules.

## Conclusion

This is the first report of sRNAs expressed in the PF stage of *S. ratti*. Most parasitic nematodes do not have genetically identical parasitic and free-living adult stages, and *S. ratti* therefore offers an almost unique opportunity to identify sRNAs specifically associated with parasitism, and to investigate sRNA-mediated targeting of TEs in a parasitic nematode. We directly compared the sRNAs expressed in the PF and FLF and key findings from this work are an identification of a novel family of 21-22U piRNA-like sRNAs in the parasitic stage of *Strongyloides* and differential expression of 27GAs across life cycle stages. These putative siRNAs originate from the X-chromosome and were predicted to target X-chromosome associated TEs. TE-targeting sRNAs were particularly evident in the PF, suggesting increased levels of TE regulatory activity associated with the parasitic life cycle stage.

## Materials and Methods

### Collection of *S. ratti* and sequencing

#### Parasite maintenance

Wistar male rats aged 4 - 6 weeks were used to maintain *S. ratti* (strain ED321) by serial passage injections of 1000 iL3 prepared by faecal culture (in 23°C for 5 days) as described by Hino *et al* (2014). All animal experiments in this study were performed under the applicable laws and guidelines for the care and use of laboratory animals, as specified in the Fundamental Guidelines for Proper Conduct of Animal Experiment and Related Activities in Academic Research Institutions under the jurisdiction of the Ministry of Education, Culture, Sports, Science and Technology, Japan, 2006.

#### Collection of PF and FLF stages

Approximately 3000 iL3s in PBS were injected subcutaneously into rats. To collect PF, rats were sacrificed on 6 days post infection (dpi) and small intestines were collected. The small intestines were developed longitudinally, washed twice with prewarmed (37°C) PBS and incubated in PBS at 37 °C for 2 hours to isolate PF. PF were washed with PBS before proceeding RNA extraction. To obtain FLF, faeces collected from infected parasites (8-10 dpi) was incubated at 23°C for 3 days using 2% agar plates. PF and FLF were transferred individually to a new tube containing TRIzol (ThermoFisher) using a needle picker after quick wash by PBS. The worm samples were snap frozen in liquid nitrogen and stored at −80°C until required.

#### Extraction of RNA and library preparation

Total RNA was extracted from ~100 PF or ~50 FLF using TRIzol, according to the manufacturer’s instructions (ThermoFisher). Small RNA libraries were constructed from 50 ng of total RNA using QIAseq miRNA Library Kit (Qiagen) according to manufacturer’s instruction. For the 5’ independent-phosphate library construction, the QIAseq miRNA Library construction protocol was modified to include RNA 5’ Pyrophosphohydrolase treatment (RppH) to remove 5’ phosphates. Briefly, 50 ng of total RNA was processed up to 3’ adapter ligation step according to the manufacturer’s instruction (Qiagen). The adapter-ligated RNA was treated with 5U of RppH (New England Biolabs) at 37°C for 30 min followed by heat inactivation at 65°C for 20 min, incubated at 4°C for 5 min and proceed immediately to the 5’ ligation protocol. Sequencing was carried out using Illumina MiSeq with MiSeq reagent kit v3-150 (Illumina). A summary of samples sequenced and total read counts for each sample is summarised in **Supplementary Data 15**.

### Identification and analysis of sRNAs

#### Processing of raw reads

Fastq files were trimmed to remove adaptor sequences using umi-tools^56^.

#### Mapping of reads to the genome

The *S. ratti* genome (bioproject PRJEB125) was downloaded from WormBase ParaSite and trimmed reads were mapped to the *S. ratti* genome with BBtools ^57^ using default settings.

#### Sequence length distribution

Sequences were filtered based on their length of 18 - 30nt using the Next generation library (NGS) Toolbox perl script TBr2_length-filter.pl ^58^.

#### Identification of microRNA sequences

miRDeep2 ^32^ was used to identify known and novel miRNA sequences. Trimmed read data from all replicates and samples generated from the monophosphate-enriched library was combined and run with miRDeep2 to identify miRNAs using default settings and using known precursor and mature reference *S. ratti* sequences as references downloaded from the miRBase database (106 precursor sequences and 208 mature sequences based on ^59,7^). Mature miRNA sequences available in the miRBase database (Release 21) for ten other nematode species (*Ascaris suum, Brugia malayi, Caenorhabditis brenneri, Caenorhabditis briggsae, Caenorhabditis elegans, Caenorhabditis remanei*, *Haemonchus contortus, Heligmosomoides polygyrus, Pristionchus pacificus, Pangarellus redivivus*) used as input for mature sequences from related species. miRNA sequences were quantified using the quantifier.pl script from miRDeep2. An estimated signal-to-noise value of 10 was used as a cut-off value and only miRNAs scoring above this threshold were used for further analysis. The resulting list of miRNA sequences was used for classification of miRNA sequences, described below.

#### Annotation of TE sequences

To identify the TE sequences, we constructed de-novo repeat library for *S. ratti* using RepeatModeler2 and then passed the library RepeatMasker 4.1.1. ^60^. LTR retrotransposons were further annotated by LTR_harvest ^61^ and LTR_digest ^62^.

#### mRNA and TE expression

mRNAseq expression data for the PF and FLF was obtained from Hunt *et al* (2016). To assess TE expression, RNAseq reads were aligned to the *S. ratti* genome using STAR ^63^ with options (--outFilterMultiMapNmax 10 --winAnchorMultiMapNmax 50). TEcounts ^64^ then ran in *multi* mode to generate counts for TEs while optimising for multimapping events. Differentially expressed TEs were identified using edgeR with thresholds of count per million (CPM) > 1 in at least 2 samples, FDR < 0.01 and fold change > 2.

#### Classification of sRNA sequences

The Unitas script (version 1.7.0) ^65^ was customised to use for a non-model organism (using the -species x parameter and custom reference databases), to classify sRNA sequences. The protein-coding genes (CDS and introns were separated), rRNA and tRNA sequences for *S. ratti* (downloaded from WormBase version WS277), TE sequences and *S. ratti* miRNA sequences (see above) were used as reference databases (using the option -refseq). Sequences are first filtered to remove low complexity reads and are then identified as miRNAs based on mature and precursor reference sequences. Sequences that are not identified as miRNAs are aligned to other reference categories. At this step if a sequence aligns in more than one reference category, they are labelled as multi-mapped. Sequences that were not classified into any of the categories outlined above were assigned as sRNAs from intergenic regions. First 5’ nucleotide data for sRNA sequences was also obtained from Unitas.

#### Classification of sRNAs by length and 5’ nucleotide

The NGS_Toolbox perl script TBr2_length-filter.pl ^58^ was used to separate sRNA reads by size ^66^. To ensure that there were no repeated sequence reads between the two libraries, sequences that were found in both the 5’ pN and 5’ modification-independent libraries were removed from the 5’ modification-independent data to generate a set of sequences that had a unique 5’ polyphosphate or capped modification. Seqkit (version 0.13.2) ^67^ was used to find duplicated sequences between the fasta files.

#### Differential expression of sRNA

The edgeR (Bioconductor version 3.11) R package (version 4.0.2) ^28^ was used to carry out a differential expression analysis. Only reads with more than 2 CPM in at least two samples were retained. Significant values include only those differentially expressed sRNAs with an FDR of < 0.01 and a fold change of > 2.

#### sRNA clusters on chromosomes

To identify clusters of sRNAs in the genome, sRNAs were mapped to the genome using Bowtie2 version 2.4.1 ^68^. The sam output file was converted to bam using Samtools (version 2.1) ^69^ followed by Bedtools (version 2.28.0) ^70^ to obtain a bed file.

#### Target site prediction

To predict potential target sequences of putative siRNAs, only reads that were found in both replicates were used. The reverse complement of sRNAs was first identified using seqkit ^67^. Sequences were then mapped to the *S. ratti* coding sequence (CDS), 250 nt upstream (predicted 5’UTR region) and 500 nt downstream (3’UTR) from the CDS sequence obtained from WormBase ParaSite, using Bowtie2 (version 2.4.1), allowing for up to one mismatches *-N 0* and *--norc* to prevent alignment of the reverse complement of the sequence. Reads were also mapped using the same method to TE sequences.

#### Protein family prediction

Predicted function of the proteins that genes coded to was updated using BLAST ^71^ and InterProScan ^72^ (Supplementary Data 16).

#### Gene Ontology

topGO (version 2.42.0) ^73^ on R was used for GO enrichment analysis. GO terms associated with each gene were obtained from WormBase parasite. REVIGO ^74^ was used to cluster GO terms according to their GO function and the REVIGO output script was customised to also report the number of sRNAs that targeted each GO term.

#### Sequence logo

To identify the nucleotide richness and the presence of any conserved motifs, WebLogo ^75^ was used to find sequence motif logos. To identify upstream sequence motifs, 21-22Us were first mapped to the *S. ratti* reference genome using bowtie2 ^68^. Flanking sequences were extracted using Bedtools.

#### Dicer-signature analysis

The stepRNA tool (version 1.0.3) (https://pypi.org/project/stepRNA/) was used to search for Dicer-processing signatures using default settings. Identical sRNA sequences were collapsed and used as input.

#### Same strand analysis

sRNA sequences were aligned to the genome using Bowtie2 with multimapping enabled (-f -a -N 0 --no-1mm-upfront --norc). Identification of overlapping sequences was achieved with Bedtools intersect ^70^ using *-s* and *-wo* parameters.

#### Statistical analyses

All statistical analyses were carried out in R studio version 3.6.3 ^76^.

#### Microscopy

Worms were fixed in 3% PFA for 20 minutes and washed three times in PBS. Specimen of worms were then transferred to glass-bottom dishes and imaged on a Leica Dmi8 inverted widefield microscope with a 10x objective and DIC prisms in the light-path. Z-stacks at different overlapping field-of-views covering the whole worm were acquired. We extracted the individual optical sections of each Z-stack as full resolution TIFF files and focus-merged them in Affinity Photo 1.9.2 (Serif Europe Ltd.).

## Declarations

### Ethics approval and consent to participate

All animal experiments in this study were performed under the applicable laws and guidelines for the care and use of laboratory animals, as specified in the Fundamental Guidelines for Proper Conduct of Animal Experiment and Related Activities in Academic Research Institutions under the jurisdiction of the Ministry of Education, Culture, Sports, Science and Technology, Japan, 2006.

### Consent for publication

Not applicable.

### Availability of data and materials

All sequence data from the genome projects have been deposited at DDBJ/ENA/GenBank under BioProject accession PRJDB13088. All relevant data are available from the authors.

### Authors’ Contributions

M.V., V.L.H. and T.K. conceived the study. M.S. V.L.H. and T.K. wrote the manuscript with inputs from others. A.K. and A.Y. prepared biological samples and performed sequencing. M.S., M.D., B.M., B.P., performed informatics analyses.

## Acknowledgements

Genome data analyses were partly performed using the DDBJ supercomputer system. We thank Simo Sun for assistance and comments.

## Author Information

Correspondence and requests for materials should be addressed to T.K. or V.H.

