## Supplementary Figs for "piRNA-like small RNAs target transposable elements in a Clade IV parasitic nematode"

**Contents Page:**

1. Supplementary Fig. 1. Expression of 5’ monophosphate (Np) tRNA-derived siRNAs in the PF and FLF.
2. Supplementary Fig. 2. Manual inspection of TE annotation shows confidence in the classification.
3. Supplementary Fig. 3. The proportion of TEs in *S. ratti* show an abundance of unclassified TEs present in the genome.
4. Supplementary Fig. 4. Expression of TEs in FLF and PF reveals a subset of unknown TEs that are highly expressed in the parasitic life cycle stage.
5. Supplementary Fig. 5. Distribution of 21-22Us across the genome.
6. Supplementary Fig. 6. Distribution of TEs across the genome.
7. Supplementary Fig. 7. No evidence of a ping-pong signature was observed for 21-22Us.
8. Supplementary Fig. 8. 21-22U and 27GA sRNA targeting may control TE expression in the PF.
9. Supplementary Fig. 9. Distribution of TEs targeted by significantly upregulated 27AGs.
10. Supplementary Fig. 10. TEs targeted by 27GA sRNAs that are upregulated in the PF and FLF do not appear to change expression.
11. Supplementary Fig. 11. Identification for the presence of an overlap signature in the PF-upregulated and FLF-upregulated 27AGs.

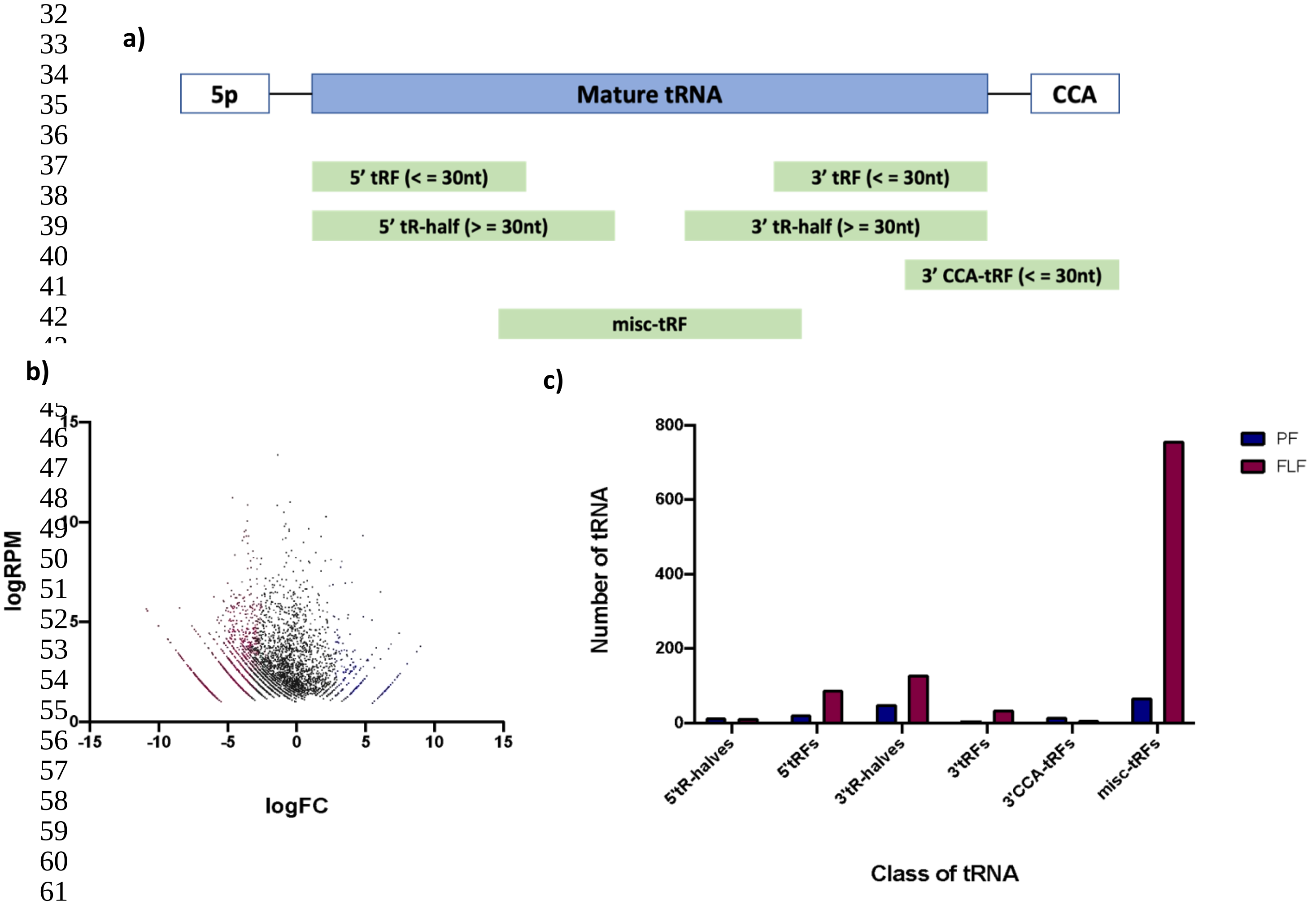


**Supplementary Fig. 1. Expression of 5’ monophosphate (Np) tRNA-derived siRNAs in the PF and FLF.** (**a**) Schematic representation of a mature tRNA containing a 5’monophosphte and a CCA tail and classification of tRNA fragments (tRF). Unitas was used to classify mature tRNA into 5’ and 3’ tRFs (tRF sequences under 30 nucleotides (nt) in length matching the end of the 5’ and 3’ of mature tRNA), 5’ and 3’ tR-halves (tRF sequences over 30 nt in length matching the end of the 5’ and 3’ of mature tRNA), 3’ CCA-tRF (tRF sequences under 30 nucleotides nt in length containing a CCA at the 3’ end) and misc-tRF, also known as internal tRFs (tRF that align mature tRNA but don’t match the 5’ or 3’ end). (**b**) Differential expression of tRNA sequences using edgeR. Significantly upregulated sequences are highlighted in pink (FLF) and blue (PF) (EdgeR Fisher’s Exact Test, FDR of < 0.01, fold change of > 2) and sequences that are not differentially expressed are shown in black (logRPM = log reads per million, logFC = log fold change). (**c**) Classification of all tRNA sequences in the PF (n = 185) and FLF (n = 1036) into tRFs.

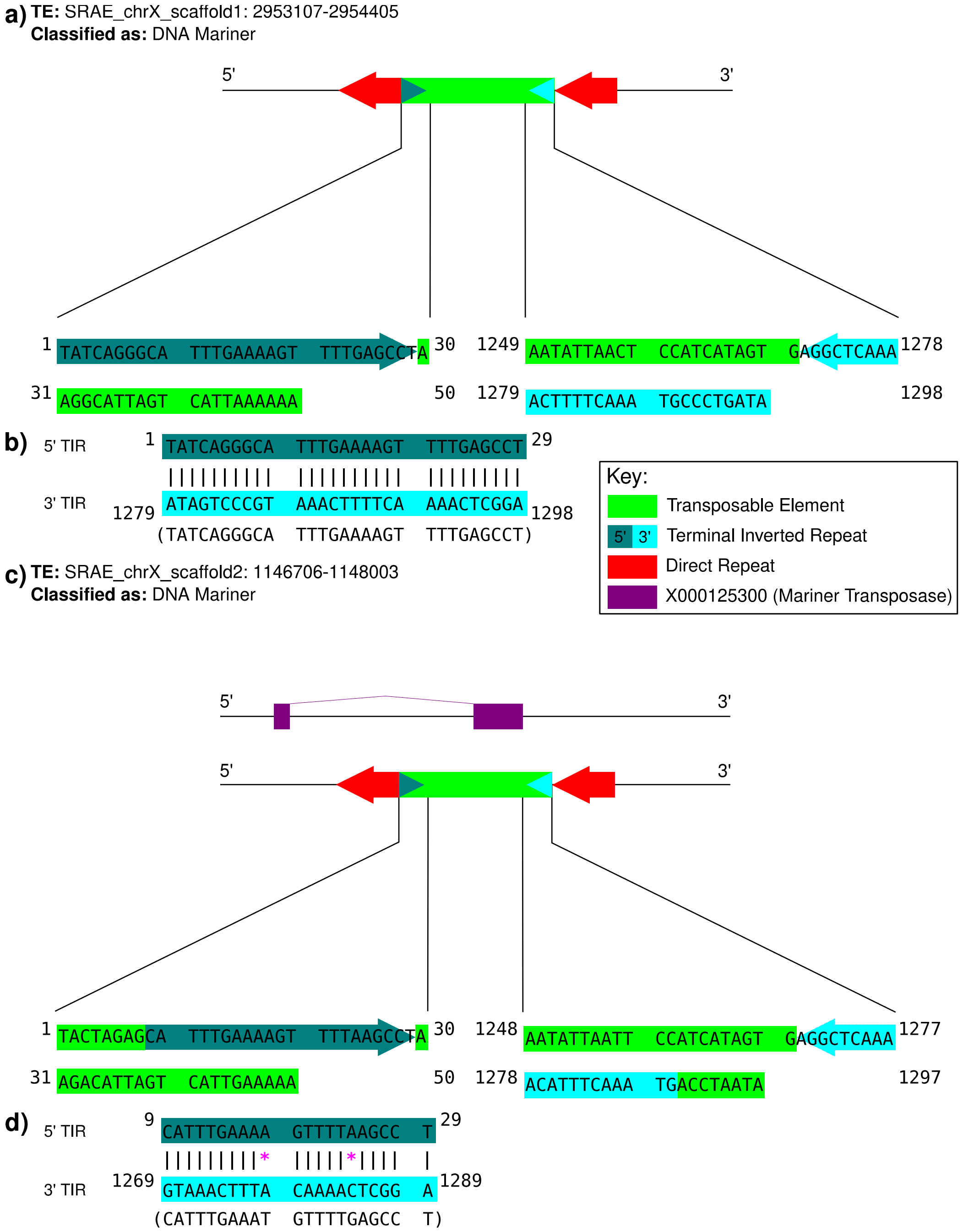


**Supplementary Fig. 2. Manual inspection of TE annotation shows confidence in the classification. (a-b)** The 50bp at the 5' and 3' end of the TE for SRAE_chrX_scaffold1_2953108_2954407 and (**b-c**) SRAE_chrX_scaffold2_1146706_1148003 extracted from the TE (green) show the 5' and 3' Terminal Inverted Repeats (TIR), turquoise and light blue respectively, confirm that the DNA Mariner classification is correct. The structure of a DNA transposon showing the 50bp region that the sequence, along with the position of the extracted sequences is shown in (**a**) and (**c**). The 5' and 3' TIR are directly compared (**b-d**) to show the reverse complementary region that is expected. The reverse complement of the 3' TIR sequence is shown in brackets below. Reverse complement matches are shown by a vertical line and mismatches by pink asterisks. For (**c**), the CDS region of a Mariner Transposase (purple) is also shown.

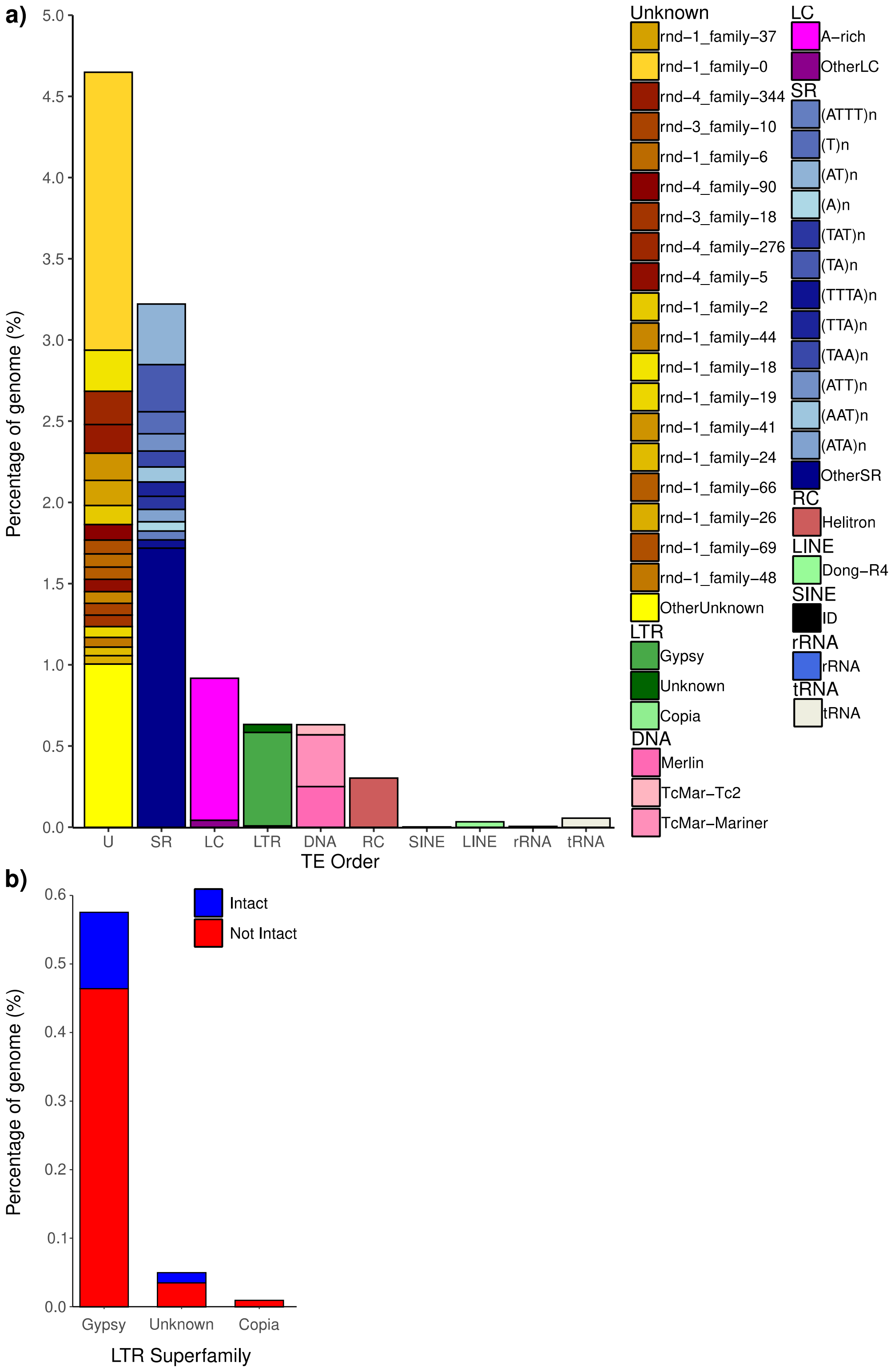


**Supplementary Fig. 3. The proportion of TEs in *S. ratti* show an abundance of unclassified TEs present in the genome.** (**a**) Barplot showing the percentage of the *S. ratti* genome for different TE orders. Bars are coloured by superfamily classification, where the individual superfamilies occupy at least 0.05% of the genome before being assigned as other. (**b**) Barplot showing the percentage of intact (blue) or non-intact (red) LTRs found by LTR harvester, in the *S. ratti* genome.

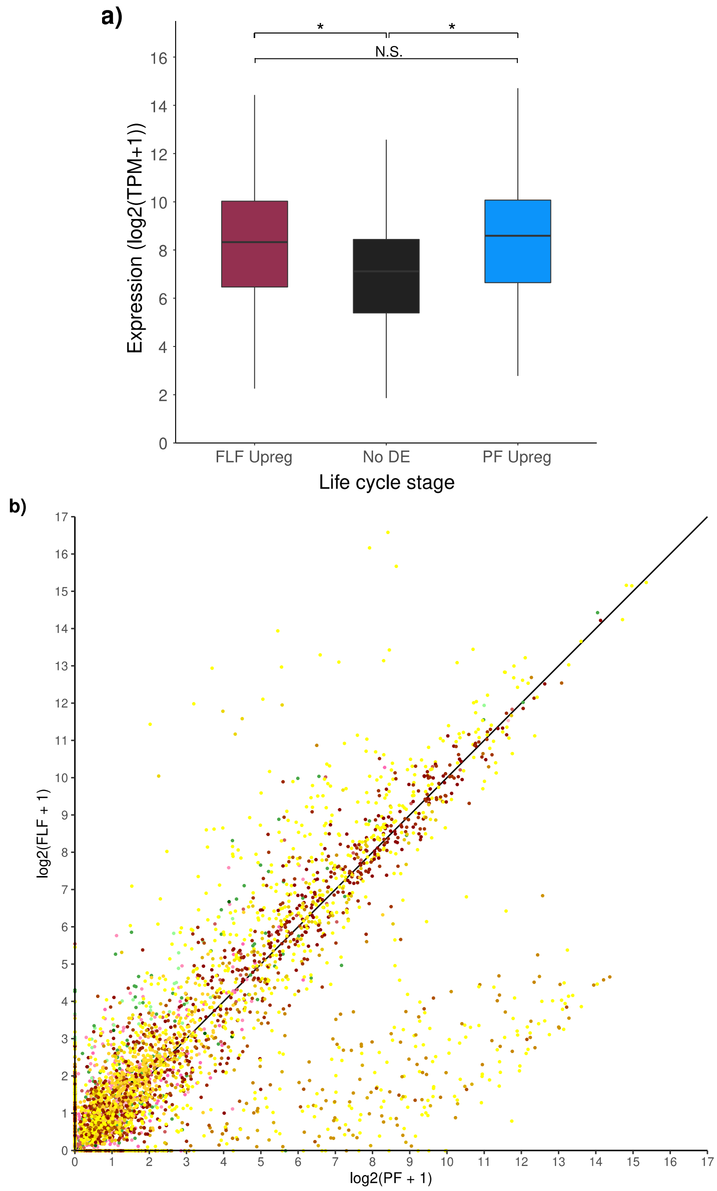


**Supplementary Fig. 4. Expression of TEs in FLF and PF reveals a subset of unknown TEs that are highly expressed in the parasitic life cycle stage.** (**a**) Boxplots showing the expression of TEs that are upregulated in PF or FLF along with those that are not differentially expressed (No DE). Transcripts per million (TPM) was used to normalise TE expression. (**b**) Scatterplot of expression of individual TEs in *S. ratti* PF (log2[TPM + 1]) compared to FLF (log2[TPM + 1]). Line shows x = y. Significance is shown by: * ($p < 0.05$). Bars are coloured by superfamily classification as supplementary **Figure S3a**.

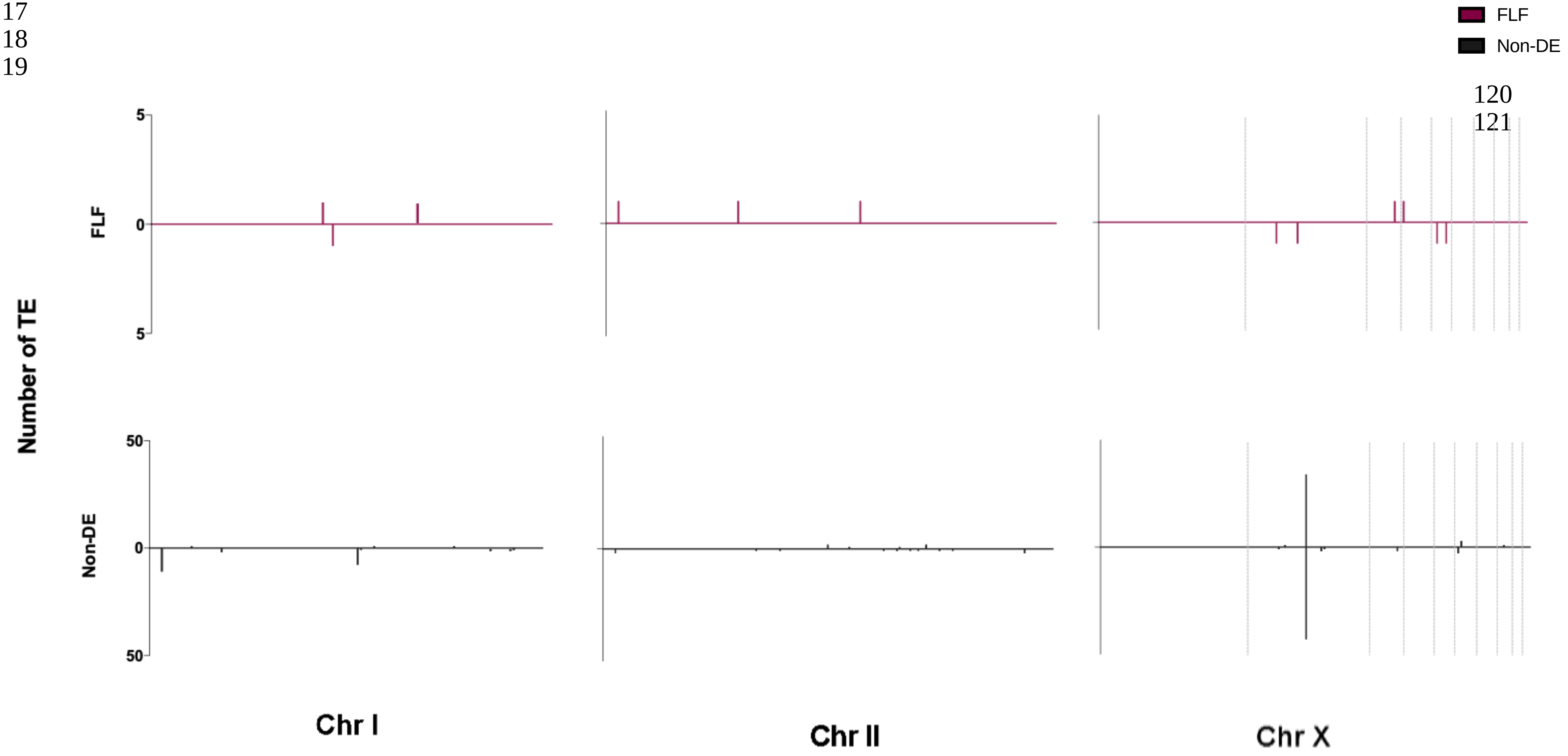


**Supplementary Fig. 5. Distribution of 21-22Us across the genome**. Expression of 21-22Us in FLF (n = 14) and non-DE (n= 195) within the genome of *S. ratti* distributed across the forward and reverse strands of the two autosomes in single scaffolds (chromosomes I and II) and the X chromosome, made up of 10 scaffolds.

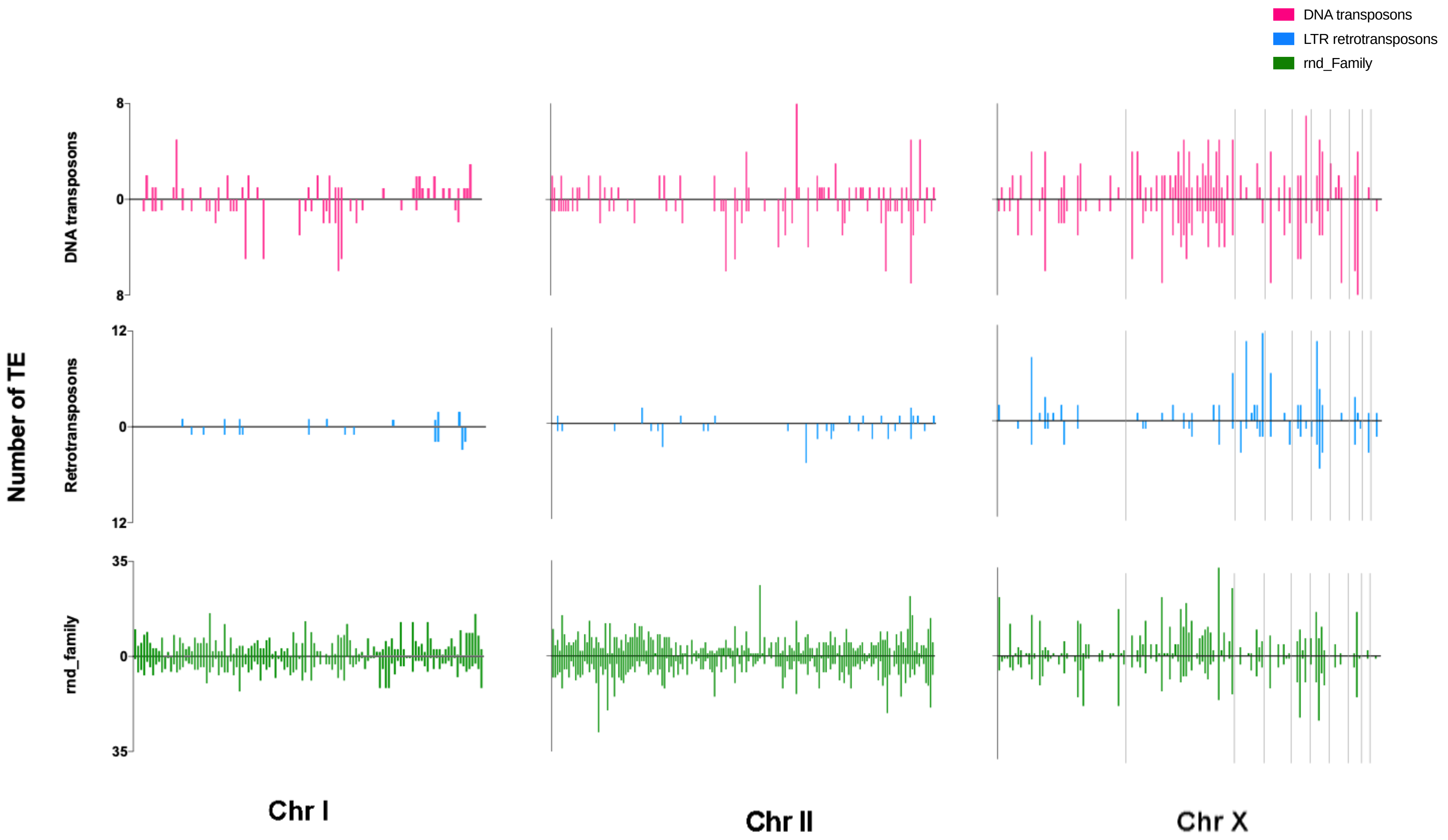


**Supplementary Fig. 6. Distribution of TEs across the genome**. Expressed TEs within the genome of *S. ratti* were identified (**SI Figure 2**) and distributed across the forward and reverse strands of the two autosomes in single scaffolds (chromosomes I and II) and the X chromosome, made up of 10 scaffolds. TE sequences were split into class I retrotransposons (n = 236) (blue), class II DNA transposons (n = 568) (pink) and the rnd_family of TEs (**see SI Table 7**) (n = 3535) (green).

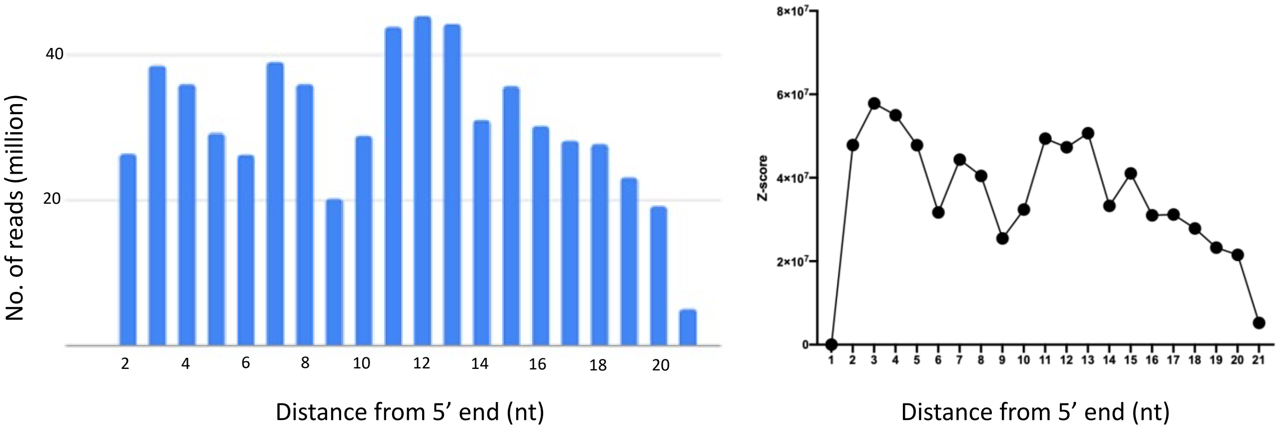


**Supplementary Fig. 7. No evidence of a ping-pong signature was observed for 21-22Us.** Unitas [66] was used to search for (a) 5’ overlaps of 21-22Us (n=1887 sequences) mapped sequence reads and (b) calculate a Z-score for the enrichment of 10 bp overlaps.

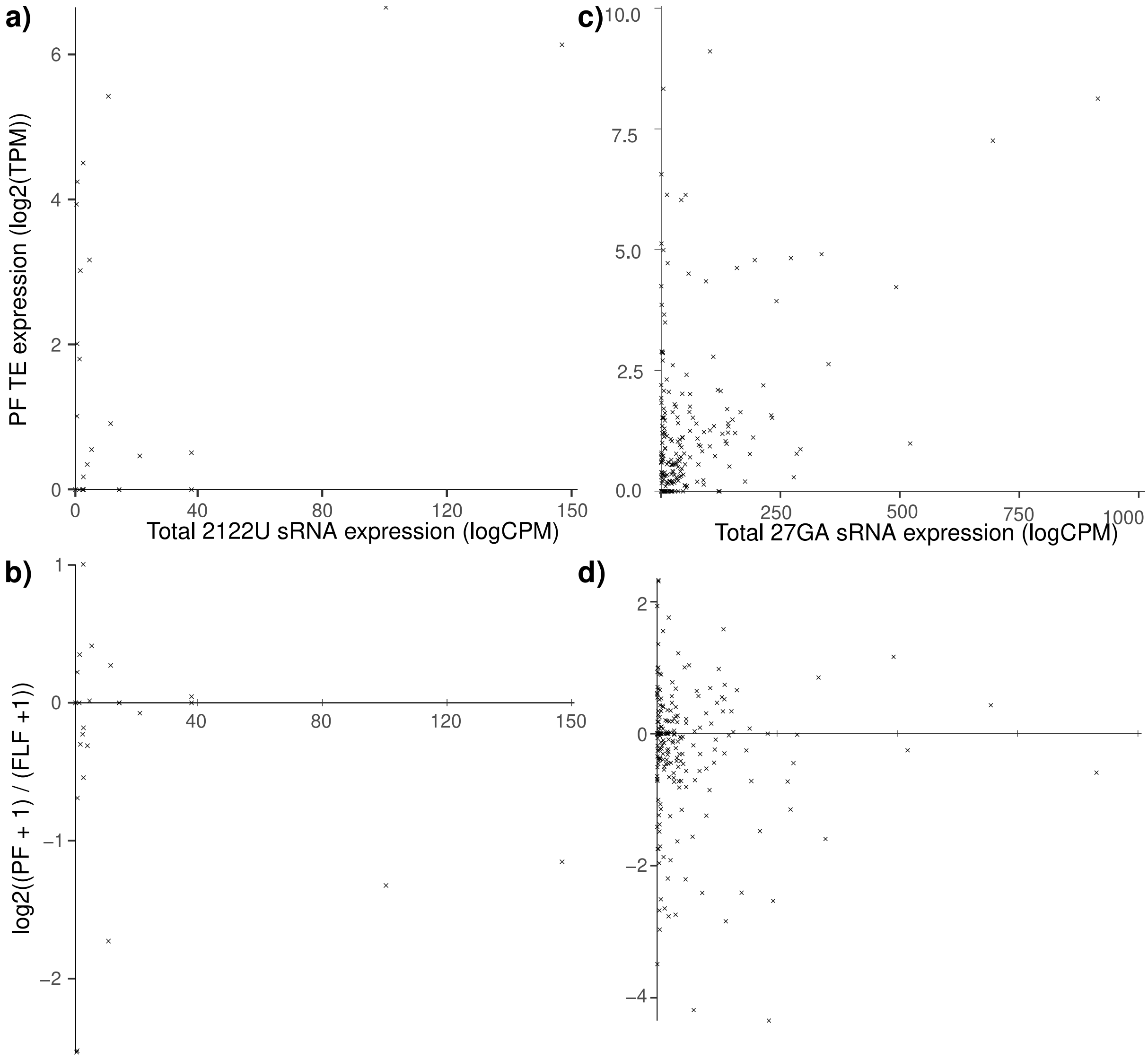


**Supplementary Fig. 8. 21-22U and 27GA sRNA targeting may control TE expression in the PF.** Scatterplots comparing total siRNA expression (logCPM) against (**a**)(**c**) PF TE expression (log2[TPM]) only or (**b**)(**d**) the difference in expression between PF and FLF, log2[RPF/FLF], for each TE that is targeted by a 2122U sRNA (**a**)(**b**) or 27GA sRNA (**c**)(**d**). TE expression was normalised using Transcripts per million (TPM). Ratios (RPF/FLF) are the same as supplementary **Figure S3**.

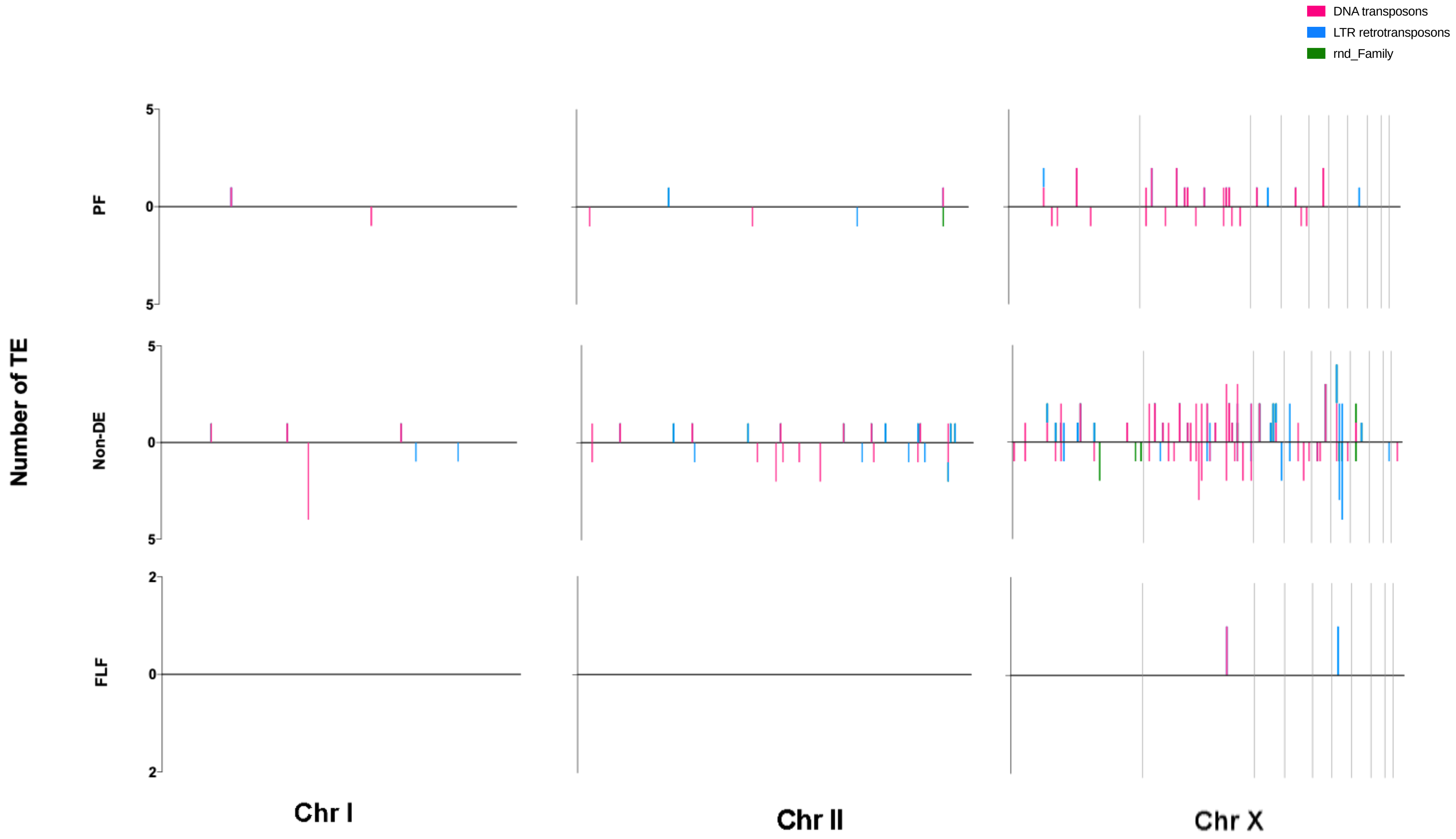


**Supplementary Fig. 9. Distribution of TEs targeted by significantly upregulated 27AGs.** TE targets of 27AGs in the PF (n = 39), FLF (n = 2) and non-DE (n = 171) were mapped and distributed across chromosome I, II and X chromosome split into 10 scaffolds. TE were split into class I retrotransposons (blue), class II DNA transposons (pink) and the rnd_family of TEs (green).

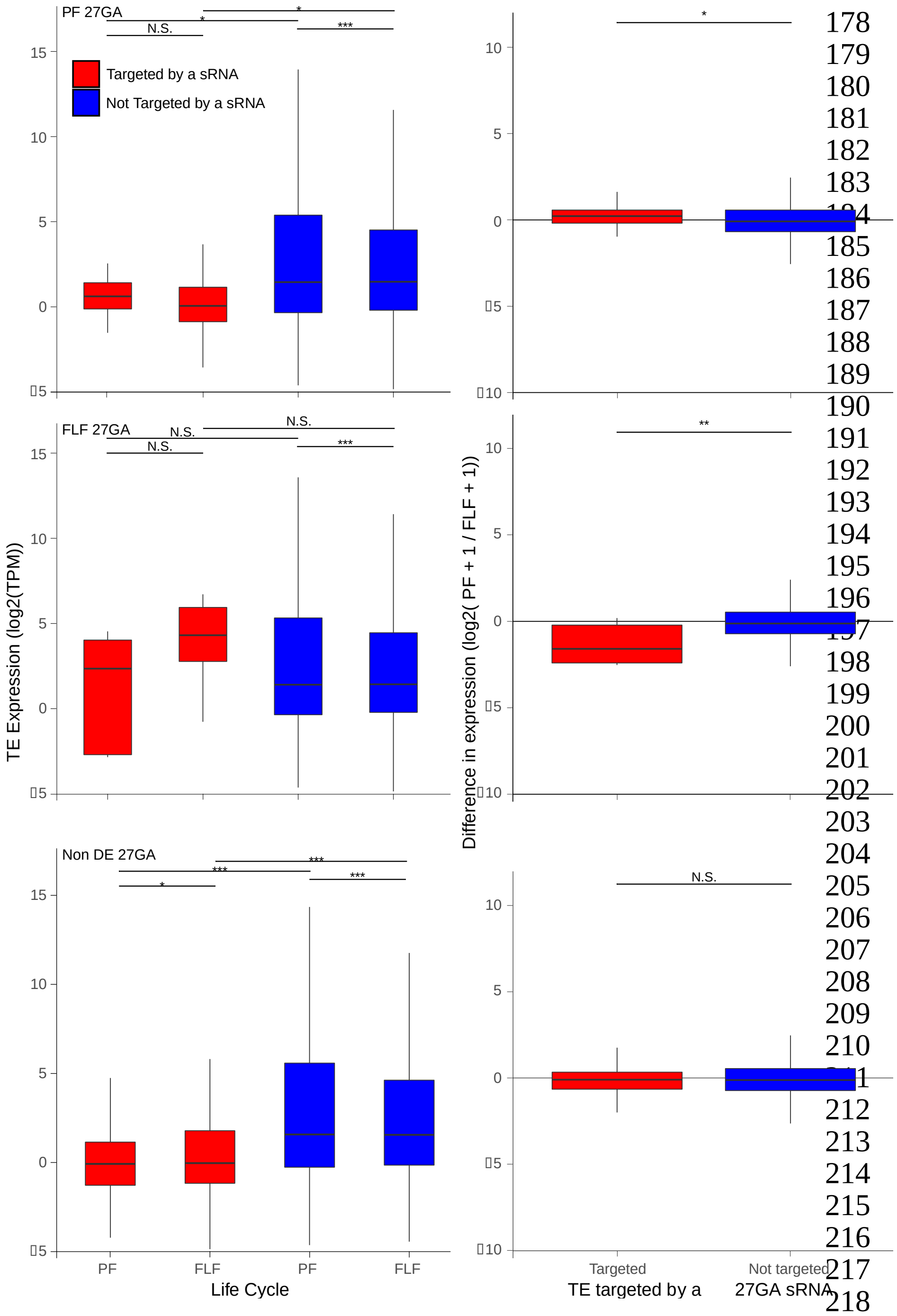


**Supplementary Fig. 10. TEs targeted by 27GA sRNAs that are upregulated in the PF and FLF do not appear to change expression.** (**a-c**) Boxplots showing the difference in expression (log2[TPM]) between TEs targeted (red) or not targeted (blue) in the PF and FLF life cycle stages for PF upregulated (**a**), FLF upregulated (**b**) and non-DE expressed (**c**) 27GA sRNAs. (**d-f**) Boxplots show the ratio of expression of TEs, between PF and FLF, that are targeted or not targeted by a 27GA sRNA that is upregulated in the PF (**d**) or FLF (**e**) life cycle stages or not -DE expressed (**f**). Ratios (PF/FLF) are the same as **Figure S4**. Dots show the log2[PF/FLF] for individual TEs. The boxes show Q1, median and Q3. TE expression was normalised using Transcripts per million (TPM). Significance is shown by: * ($p < 0.05$), ** ($p < 0.01$) and *** ($p < 0.001$).

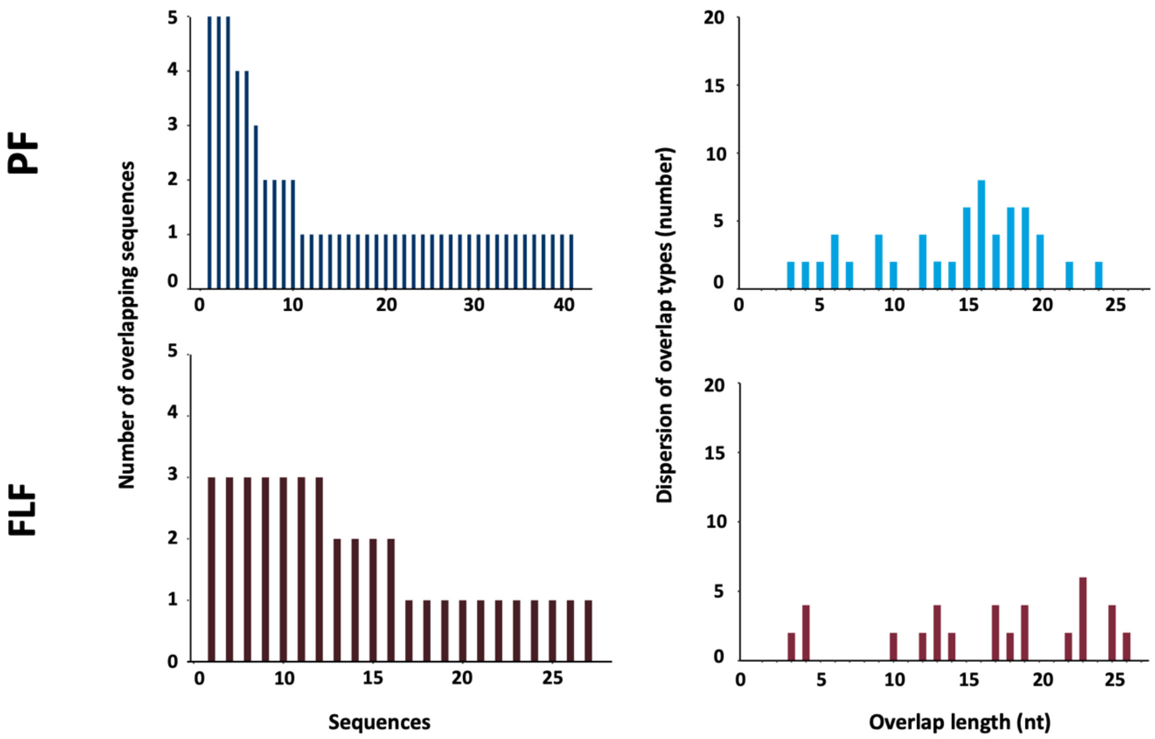


**Supplementary Fig. 11. Identification for the presence of an overlap signature in the PF-upregulated and FLF-upregulated 27AGs.** Left figure shows the number 27AGs that overlap with other 27GAs in the PF (20.2%) and FLF (42.3%). Right figure showing the overlap lengths of siRNAs against the 27AGs.
